# Origins of *de novo* chromosome rearrangements unveiled by coupled imaging and genomics

**DOI:** 10.1101/2024.08.15.607890

**Authors:** Marco Raffaele Cosenza, Alice Gaiatto, Büşra Erarslan Uysal, Álvaro Andrades Delgado, Nina Luisa Sautter, Michael Adrian Jendrusch, Sonia Zumalave Duro, Tobias Rausch, Aliaksandr Halavatyi, Eva-Maria Geissen, Patrick Hasenfeld, Isidro Cortes-Ciriano, Andreas Kulozik, Rainer Pepperkok, Jan O. Korbel

**Affiliations:** Genome Biology Unit, European Molecular Biology Laboratory (EMBL), Heidelberg, Germany; Molecular Medicine Partnership Unit (MMPU), EMBL, University of Heidelberg, Heidelberg, Germany; Department of Pediatric Oncology, Hematology, and Immunology, University of Heidelberg and Hopp Children’s Cancer Center, Heidelberg, Germany; European Bioinformatics Institute (EMBL-EBI), Hinxton, Cambridgeshire, UK; Core Facilities Unit, EMBL, Heidelberg, Germany; Data Science Centre, EMBL, Heidelberg, Germany; Cell Biology and Biophysics Unit, EMBL, Heidelberg, Germany; Bridging Research Division on Mechanisms of Genomic Variation and Data Science, German Cancer Research Center (DKFZ), Heidelberg, Germany

## Abstract

Chromosomal instability results in widespread structural and numerical chromosomal abnormalities (CAs) during cancer evolution^1–3^. While CAs have been linked to mitotic errors resulting in the emergence of nuclear atypias^4–7^, the underlying processes and basal rates of spontaneous CA formation in human cells remain under-explored. Here we introduce machine learning-assisted genomics-and-imaging convergence (MAGIC), an autonomously operated platform that integrates automated live-cell imaging of micronucleated cells, machine learning in real-time, and single-cell genomics to investigate *de novo* CA formation at scale. Applying MAGIC to near-diploid, non-transformed cell lines, we track CA events over successive cell cycles, highlighting the common role of dicentric chromosomes as an initiating event. We determine the baseline CA rate, which approximately doubles in *TP53*-deficient cells, and show that chromosome losses arise more rapidly than gains. The targeted induction of DNA double-strand breaks along chromosomes triggers distinct CA processes, revealing stable isochromosomes, amplification and coordinated segregation of isoacentric segments in multiples of two, and complex CA outcomes, depending on the break location. Our data contrast *de novo* CA spectra from somatic mutational landscapes after selection occurred. The large-scale experimentation enabled by MAGIC provides insights into *de novo* CA formation, paving the way to unravel fundamental determinants of chromosome instability.

## Introduction

Cancer whole genome sequencing (WGS) studies have underscored the roles of CAs in shaping mutational landscapes^8,9^. CA driver alterations outnumber base substitution drivers in pan-cancer studies^3,8^, and the genomic burden of CAs resulting in copy-number imbalances is linked to cancer recurrence and patient death^2,10^. The emergence of CAs in a cell is presumed to represent a pivotal event in tumorigenesis, which may trigger episodic or sustained chromosomal instability enhancing karyotype diversity and intra-tumor heterogeneity^2,3,6,11^. These alterations can facilitate adaptation to selective pressures during cancer progression and therapy, resulting in adverse clinical outcomes^2,3,12^.

A variety of CA formation processes have been reported, leading to numerical chromosome abnormalities, simpler interstitial structural variants (SVs) including deletions and duplications, as well as complex CAs with numerous DNA breakpoints resulting from events such as the breakage-fusion-bridge (BFB) cycle or chromothripsis^13–19^. However, unlike for base substitutions^20,21^, the relative contributions of *de novo* CA processes, as well as the baseline rate at which CAs emerge, are poorly understood. Bulk tissue sequencing of tumour genomes has shed light into the classes of DNA rearrangements observed in primary cancer^15,16,21,22^, and highlighted the role of defective DNA replication in generating small to intermediate-sized SVs^16^. Yet, as an important limitation, newly arising CAs, which are presumed to be mostly detrimental to cells and thus quickly eliminated from a cell population, are not detectable using bulk WGS^3,23^. As a result, the impact of sporadic CA formation processes on the somatic mutational landscape is under-explored.

To comprehensively investigate CA formation, it is imperative to examine CAs in single cells^3^ – sampled before selective forces can act. Recent reports have established that a single DNA lesion can trigger a cascade of alterations leading to complex CA formation^4,6,17^. Mitotic errors serve as intermediate steps for these cascades, resulting in atypical nuclear structures (nuclear atypias), such as micronuclei and chromatin strings^3,24^. Live-cell microscopy combined with (semi)-manual cell selection and single-cell sequencing, has been utilised to elucidate roles of nuclear atypias in the formation of complex CAs, revealing insights on the mechanisms underlying chromothripsis^4–7,17,25,26^. Yet, these approaches are labour-intensive and have examined only a relatively small number of single-cell genomes per study, with a particular focus on CAs that have been induced by biochemical or genetic techniques. This has limited insights into the spectrum of CAs that spontaneously arise in cells, leaving gaps in our understanding of fundamental chromosomal instability processes.

To address these limitations, we devised <u>M</u>achine-learning <u>A</u>ssisted <u>G</u>enomics- and-Imaging <u>C</u>onvergence (MAGIC). This platform couples automated confocal microscopy in live cells, machine learning (ML) for real-time nuclear atypia assessment, targeted cell illumination, and cell sorting. This enables the imaging-based selection and precise isolation of target cells from a heterogeneous cell population. The target cells are subsequently subjected to single-cell genomics or phenotype analysis, enabling systematic study of *de novo* CA at large-scale. Using micronuclei as an imaging biomarker for cells at risk for CA formation^3,5,6,27^, we applied MAGIC to comprehensively investigate sporadic CA formation in two non-transformed cell lines. Our findings reveal highly heterogeneous CA landscapes, yet with recurrent patterns pointing to a relatively small number of processes driving their emergence. By reconstructing mitotic histories, we highlight dicentric chromosomes as a major contributor to CA formation. We estimate the rate of *de novo* CAs associated with different mitotic errors and evaluate the influence of *TP53* deficiency. Furthermore, by integrating MAGIC with DNA double-strand break (DSB) engineering, we uncover a crucial impact of derivative chromosome structure on CA spectra, and highlight isoacentric formation as a previously unrecognised process facilitating the rapid amplification and asymmetric segregation of large DNA segments. By enabling the convergence of imaging and genomics, MAGIC advances research on CA formation processes.

## Results

### MAGIC: analysis of spontaneous CA formation by automated imaging and single-cell genomics

#### Mitotic error profiles in unperturbed non-tumorigenic cells

To comprehensively investigate *de novo* CAs arising in a human cell, we devised MAGIC, which couples automated confocal microscopy with targeted illumination and single-cell genomics to gain insights into CA formation from studying nuclear atypias (**Fig. 1A**). To better represent CA formation landscapes during an early stage of tumorigenesis, we chose two non-tumorigenic, near-diploid cell lines that maintain a relatively stable karyotype and are widely used in chromosomal instability studies^4,5,28–30^: MCF10A cells, derived from normal breast tissue and spontaneously immortalised, and hTERT RPE-1 cells (RPE-1) of retinal pigment epithelial origin. During cell division, both cell lines occasionally form micronuclei^31–33^, the collapse of which has been causally linked to CA formation including chromothripsis events^3,5,6^.

**Figure 1:**
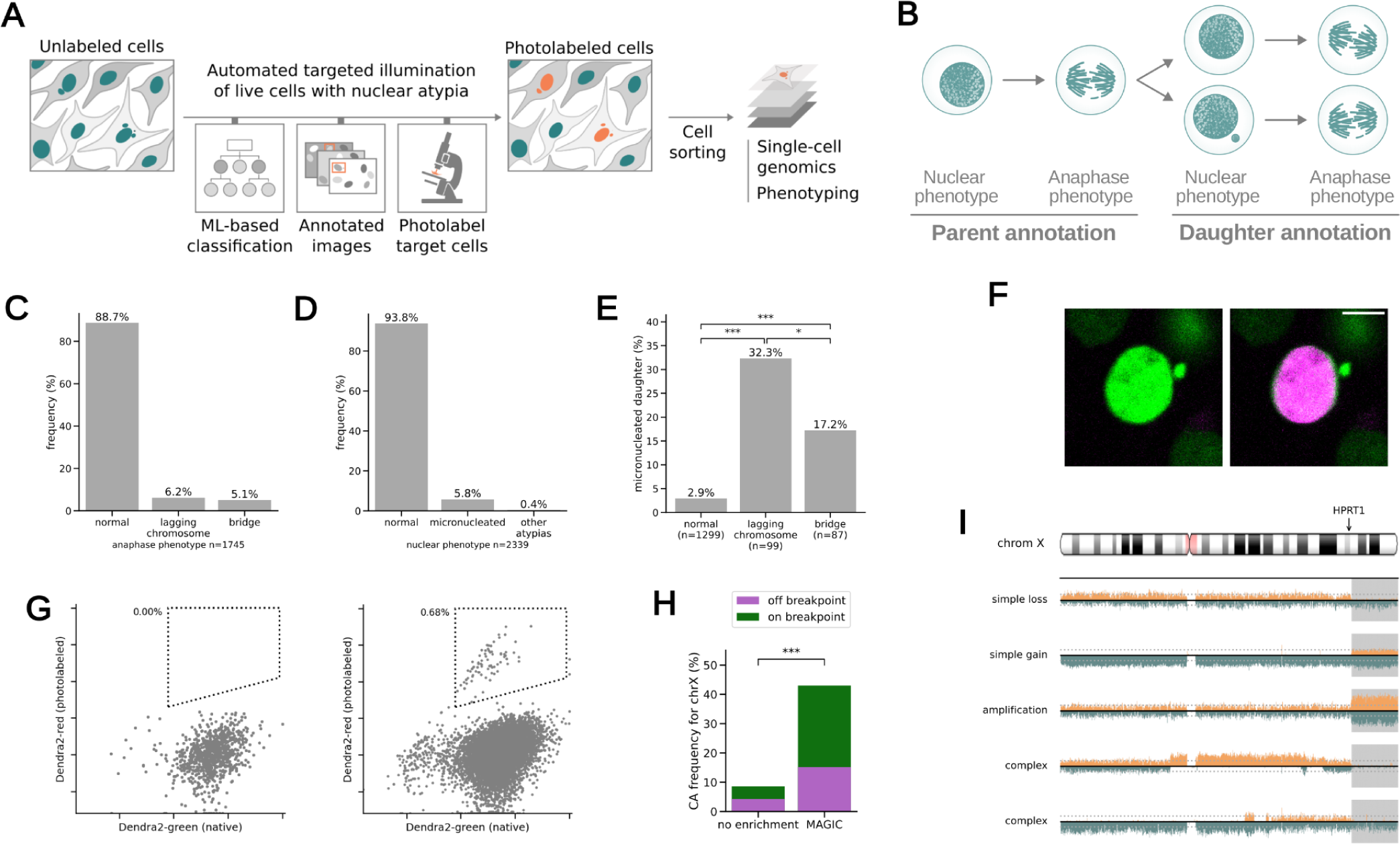
MAGIC enables the single-cell genomic characterisation of de novo CAs at large-scale. (A) Overview of MAGIC, from left to right: An automated microscope analyses a cell population with a heterogeneous morphology (i.e. micronuclei), performing targeted illumination using 3 steps: (i) a ML-model classifies cells to retrieve a probability score for each cell; (ii) areas for targeted illumination are identified; (iii) an automated laser beam selectively photolabels target cells. The steps are repeated to reach a desired yield. Lastly, photolabeled cells are isolated for downstream analysis. (B) Annotation strategy for long-term live-cell imaging of MCF10A cells. Anaphase and nuclear phenotypes were assessed over two generations, to observe at least one entire cell cycle; cells from the first annotated generation are referred to as parents, while cells from the second as daughters. A total of 821 genealogies were annotated, 765 of which generated progeny. (C,D) Overall frequency of anaphase (C) and nuclear (D) phenotypes in MCF10A cells from long-term live-cell imaging, over all genealogies, 1,750 mitoses and 1,795 nuclei were annotated, respectively. (E) Frequency of daughter micronucleation associated with different anaphase phenotypes. (F) MCF10A cells constitutively expressing H2B-Dendra2 before (left) and after (right) photolabeling via targeted-illumination by MAGIC. H2B-Dendra2 emits green fluorescence in its native state, which shifts towards red after photoconversion (here represented in magenta). Scale bar: 10 μm (G) FACS profile of MCF10A H2B-Dendra2 cells in control (left) and photolabeled (right) conditions. Native (green) versus photoconverted (red) fluorescence is plotted. The sorting gate is represented by a dashed line. (H) Frequency of CAs affecting chromosome X in single cells randomly sampled, or selected using the MAGIC platform, after targeted DBS induction. On-breakpoint CAs (green bar) refer to the HPRT1 locus. (I) Examples of CAs affecting chromosome X and involving a breakpoint at HPRT1 locus (see chromosome ideogram above). Grey background: acentric fragment created through targeted DSBs. (for significance levels see Methods, * P<0.05, **P<0.01, ***P<0.001).

To generate pilot data for setting up the MAGIC platform, we initially performed a comprehensive manual annotation of nuclear and mitotic phenotypes across two generations in MCF10A cells, characterising 821 parental cells and their progeny altogether (**Fig. 1B**). We find that sporadically arising anaphase bridges and lagging chromosomes are the predominant type of mitotic error, occurring in 5.1% and 6.2% of all mitoses, respectively (**Fig. 1B**). During interphase, 6.2% of cells display nuclear atypias, out of which micronuclei are the by far most common type (5.8%) (**Fig. 1C**). In contrast, other types of nuclear atypia, such as large nuclei indicative of whole-genome duplication, are rare, leading us to concentrate our subsequent analyses mostly on micronuclei. We find that mitotic errors result in the formation of at least one micronucleated daughter cell in 32.3% and 17.2% of cases for lagging chromosomes and chromatin bridges, respectively, underscoring that both types of mitotic error converge on micronucleation (**Fig. 1D**). Furthermore, micronucleated cells are ∼9.5 times more likely to generate a micronucleated daughter cell, compared to cells with normal (round) nuclei, with 37.7% of these daughters also bearing a micronucleus (**Fig. S1A**). Accordingly, we detect a high frequency of anaphase errors in daughters from cells showing an abnormal anaphase (**Fig. S1B**). This ‘self-propagating’ nature of micronucleated cells implies that mitotic errors can result in transient nuclear atypia formation over consecutive cell cycles, possibly triggering episodic *de novo* CAs.

We continued these systematic annotations, by examining micronucleated MCF10A cells with respect to their propensity to generate viable daughters. Micronucleated cells exhibit a significantly longer cell cycle duration compared to normal cells (**Fig. S1C**); accordingly, 22.2% undergo cell cycle arrest compared to 6.6% of normally nucleated cells (**Fig. S1D**). When considering the nuclear phenotype of the parental cell, we find that having a micronucleus delays the cell cycle also in those cases where the parental cell already carries a micronucleus (**Fig. S1E**). In summary, anaphase errors, such as lagging chromosomes and chromosomal bridges, converge in the formation of micronuclei, suggesting the latter represent an interphase biomarker for the enrichment of cells potentially undergoing CAs. While micronuclei formation can result in cell cycle delay and arrest, these consequences are neither absolute nor immediate, with relevant cell subsets eventually regaining normal nuclear morphology. We reasoned that these cellular properties would facilitate the automated isolation of viable micronucleated cells subject to *de novo* CAs using the MAGIC platform.

#### A machine learning-enabled adaptive feedback loop driving the MAGIC platform

We thus chose micronuclei as the primary imaging biomarker for developing MAGIC (**Methods**). Particularly, in an automated process, cells bearing micronuclei are systematically selected by MAGIC utilising an adaptive-feedback loop (**Fig. 1A, Fig. S2A**). This loop is driven by computational frameworks designed to steer its ML, image analysis, and photolabeling components.

In brief, a confocal image is acquired and provided for ML in real-time; if a cell of interest is identified by MAGIC, information on its location is utilised and its nucleus photolabeled with an automated laser. Then, MAGIC moves to the next image, re-initiating this loop. MAGIC operates autonomously for up to 24 hours, examining tens of thousands of cells in order to photolabel hundreds of cells exhibiting the desired rare nuclear morphology for downstream analyses.

#### Targeted photolabeling and cell sorting

The photolabeling step of MAGIC capitalises on dyes with unique characteristics. The expression of the Dendra2 protein allows tagging cells-of-interest through the stable transition of emitted fluorescence from green to red^7,34^, upon gentle and targeted illumination with a 405 nm laser. For our study, we engineered MCF10A and RPE-1 cells to stably express H2B-Dendra2, facilitating nuclear morphology visualisation and serving as a biomarker for nuclear atypias as well as a photolabeling dye (**Fig. 1F**). We find that photolabeling increases with the illumination up to a plateau (**Fig. S2B**), with the maximum fluorescence dependent on the level of H2B-Dendra2 expression (**Fig. S2C**). We fine-tuned photolabeling conditions to achieve a red fluorescence increase of ∼22-fold after illumination (**Fig. S2D**), without any detectable phototoxicity-mediated cell death within the average duration of the experiments (**Fig. S2E**). As an alternative dye, we synthesised DACT-1 (**Methods**), a small molecule previously used for cell tracking^35^, which freely diffuses into cells and is retained after cytoplasmic esterase activity removes an ester group, allowing for photo-activation under similar conditions as Dendra2 (**Fig. S2F**). Since DACT-1 requires no genetic manipulation, it enhances the versatility of MAGIC. Utilising FACS, we observe distinct populations that represent photolabeled cells with either dye (**Fig. 1G, S1G**), confirming that target cells are efficiently sorted.

#### Benchmarking and tuning the ML components of MAGIC

To enable the real-time detection of micronuclei, we trained micronuclei classifiers on manually annotated nuclear atypias (**Methods**). We find that an extreme gradient boosting-based ML framework (XGBoost)^36^ performs similarly to a deep learning-based convolutional neural network (**Fig. S2H** and **Supplementary Methods**). For the MAGIC experiments detailed below, we selected XGBoost due to its model explainability, streamlined implementation, and the fewer training examples it requires. We tuned the parameters of this ML framework to achieve a precision of at least 90% and a recall of 50%, allowing for an acceptable balance between specificity and sensitivity.

#### Targeted CRISPR/Cas9 manipulation reveals de novo Cas

Having demonstrated the effectiveness of MAGIC in cell sorting, we next explored its utility for identifying *de novo* CAs. To experimentally verify that discovered CAs originate from chromosomes entrapped in micronuclei, we generated DSBs at the *HPRT1* locus on chromosome X, by applying CRISPR/Cas9 in MCF10A cells (**Methods**). Cas9-mediated DSBs have previously been reported to result in acentric fragments^37^ that cannot participate in chromosome segregation and end up in micronuclei (**Fig. S3A**). This can result in CA formation involving the cut fragment^37^. Quantifying nuclear defects upon targeted DSB generation, we find that micronucleation is increased by ∼4.8-fold, suggesting potentially increased CA formation involving the cut site^37^.

To enable the discovery of CA classes resulting in copy-number imbalances, we coupled MAGIC with Strand-seq^38^, for single-cell template strand sequencing. Unlike other single-cell genomic techniques, Strand-seq uniquely preserves haplotype identity across an entire homolog^38,39^, which enables sensitive detection of simple and complex CA classes at intermediate sequence coverage^39,40^. We isolated cells 48h after Cas9-manipulation, giving the cells sufficient time to recover and to allow for BrdU labeling^38^ for Strand-seq library preparation. We performed single-cell genomic sequencing on 85 and 93 cells, respectively, with and without MAGIC-based selection for micronucleated cells. To perform CA discovery, we utilised single-cell tri-channel processing^39^, a computational approach integrating read-depth, strand-state, and haplotype-phase from the Strand-seq data (**Methods**). We observe a strong increase in CAs in the selected cell fraction (**Fig. S3B**), with 37 (45%) of micronucleated cells showing at least one CA on the X chromosome, corresponding to a 5-fold enrichment over nuclei isolated without enrichment. Moreover, 24/37 (64.8%) of the CAs in the enriched sample contain a breakpoint at the *HPRT1* locus (**Fig. 1H**), demonstrating that the Cas9-mediated DSBs triggered *de novo* CAs at the cut site.

When resolving each CA affecting chromosome X, we observe a vast diversity of CA classes, all presumably originating from the same cut site (**Fig. 1I, S3C**). Within the micronucleated cells, 18/37 (48.6%) of CAs show an isolated loss or gain of the cut fragment, suggesting abnormal segregation of the acentric fragment. Moreover, we find twelve CAs comprising terminal deletions from 18 to 70 Mb in size, and six are terminal duplications 22 to 76 Mb in size. We also find evidence for complex CAs in nine cases, and amplifications of the cut acentric fragment in an additional nine cases (**Fig. S3D**). Homolog-resolved analyses of the Strand-seq data shows that while CAs can affect either haplotype, in all nine cases of presumed complex CAs with unambiguous haplotype information^39^ we find that the respective rearrangements map to the same homolog (**Fig. S3E**), in support of complex CA formation. These data underscore the capability of MAGIC to isolate cells subject to *de novo* CAs.

#### Verification of de novo CA formation through analysis of sister cell pairs

Previously, the analysis of sister cells has been used to reconstruct complex CA processes, but necessitated laborious cell selection techniques^5^. We explored the utilisation of Strand-seq data to automate the detection of sister cells from large single-cell genomic datasets. Reciprocal template strand inheritance and sister chromatid exchange (SCE) events^38^ from Strand-seq offer uniquely identifiable records of sister cell relationships (**Fig. S3F**). Making use of these records, we devised a method to identify sister cell pairs through analysing single-cell genomes (**Methods**). By applying this method to our full dataset, we detect two sister cell pairs in the enriched sample, one of which exhibits a CA. Genomic analysis of the latter pair shows a cut fragment that is asymmetrically inherited in the sisters, thus independently verifying *de novo* CA formation (**Fig. S3G**).

### Landscape of spontaneous CA formation in micronucleated cells

#### Extensive spontaneous de novo CA formation in micronucleated cells

While biochemically or genetically induced micronuclei have been employed to systematically study CA formation^4–6,17^, how sporadically arising micronuclei – without experimental perturbations – may trigger distinct classes of CAs is under-explored. To address this gap, we coupled MAGIC with Strand-seq, investigating *de novo* CAs using spontaneous micronuclei as an imaging biomarker. We first focused on MCF10A, sequencing 142 single-cell genomes from micronucleated cells. By performing single-cell genome analysis (**Methods**), we identify 124 CAs with 54.9% of cells exhibiting at least one CA (**Fig. 2A**). When compared to 115 single-cell genomes from cells with a normal nucleus (‘controls’), we detect a marked enrichment of CAs (3-fold; *P*=1.75e-9; Fisher’s exact test), suggesting widespread CA formation in the micronucleated cell population. We further expanded our analysis by including RPE-1 cells, sequencing the genomes of 166 micronucleated cells and 68 controls. Unlike for MCF10A, we find a non-significant enrichment of CAs in micronucleated cells (*P*=0.69), indicating that RPE-1 cells have a more stable karyotype (**Fig. 2A**). These data imply that spontaneously micronucleated cells represent a relevant source of *de novo* CAs, although this appears to vary with the cell-line context.

**Figure 2:**
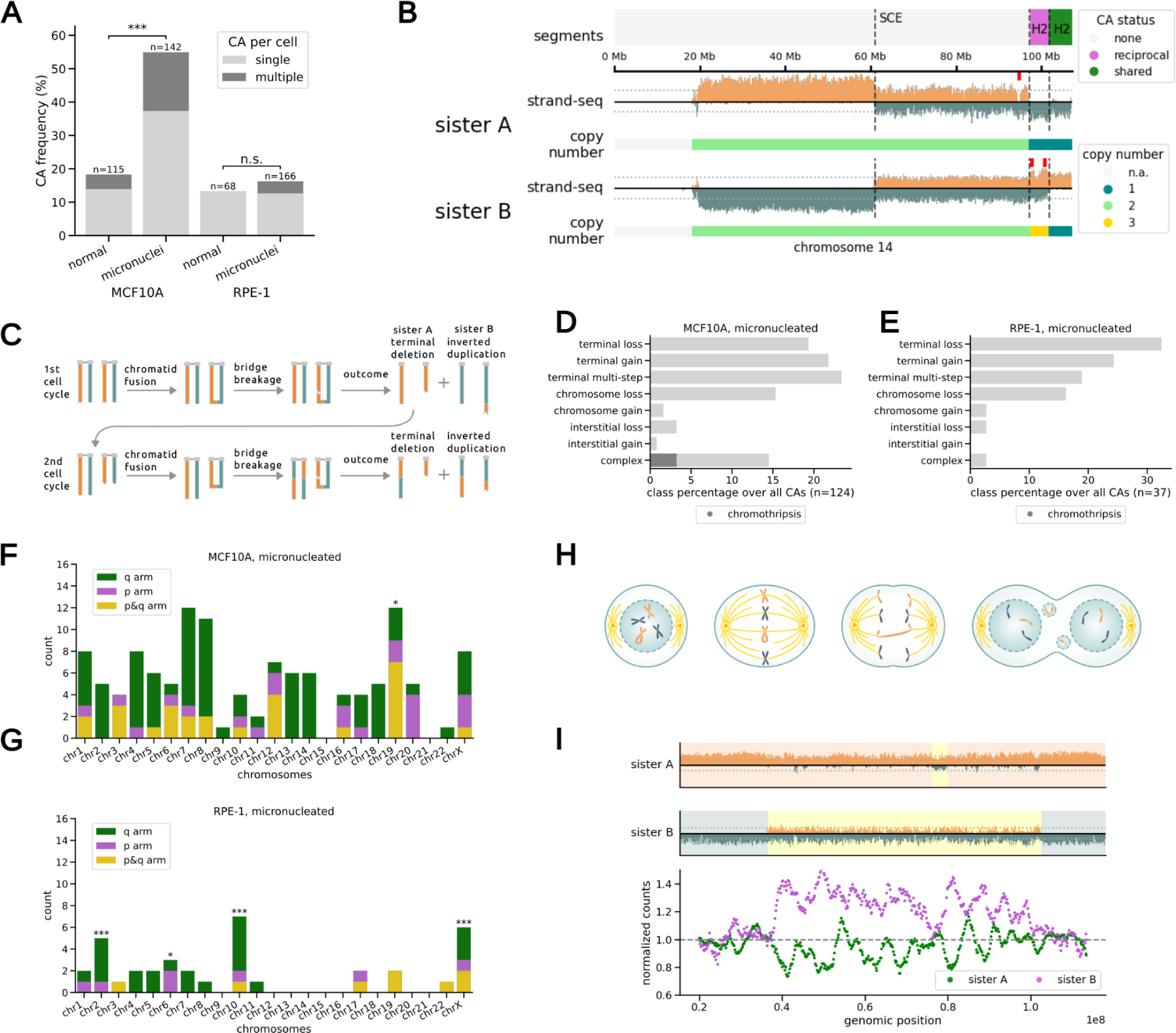
CA landscape of spontaneous micronucleated cells in near-diploid human cell lines. (A) CAs detected per cell (P-values based on Fisher’s exact test). (B) Reciprocal CAs between sister cells. Segment annotation: dashed lines indicate an SCE, and demarcate shared and reciprocal segments. Red ticks mark points of inferred complex CA formation. (C) Mitotic history reconstruction for reciprocal CAs in (B). Orange and teal colour correspond to Watson (W) and Crick (C) template strand orientations, respectively. (D,E) Breakdown of CA classes in micronucleated MCF10A (D) and RPE-1 (E) cells, class percentage over all CAs. See methods section for CA classification criteria. (F,G) CA count per chromosome in micronucleated MCF10A (F) and RPE-1 (G) cells (Binomial testing was used to identify enrichments) (H) Scheme showing mitosis with a dicentric chromosome: sister chromatid fusion generates a dicentric. During anaphase, the dicentric chromosome forms a bridge, which can rupture and potentially form micronuclei in daughter cells. (I) Chromothripsis affecting chromosome 13, with reciprocal segment inheritance into sister cells seen for a large fraction of the affected homolog. Top, Strand-seq data. Bottom, smoothened, normalised read counts along chromosomal positions.

#### Reconstructing de novo CAs arising over consecutive cell cycles

To independently corroborate CA formation in spontaneously micronucleated cells, we computationally searched for sister cell pairs among all sequenced single-cell genomes. We find 12 sister cell pairs among the 142 micronucleated MCF10A cells (**Fig. S4A**), out of which 7 (58%) show reciprocal CA segregation, verifying widespread *de novo* CA formation (**Fig. S4B**). Four sister cell pairs show no CAs, implying micronucleation does not always result in CAs. We also find three sister cell pairs with shared CAs, which in two cases are found in addition to a reciprocal CA, indicative of CA formation during an earlier cell division, with further CAs arising during the most recent cell cycle. By comparison, we find 10 sister pairs among the micronucleated RPE-1 cells (**Fig. S4A**). Of these, only two pairs exhibit reciprocal CAs (**Fig. S4B**), while one pair harbours a shared CA, consistent with RPE-1 cells exhibiting a much lower rate of spontaneous CAs. These data imply that while sister cell pair analyses are not necessary for the discovery of *de novo* CAs with Strand-seq, they can help resolve historical rearrangements happening during prior cell divisions.

To more systematically explore CA events happening across consecutive cell cycles, we conducted an in depth analysis of all seven MCF10A sister cell pairs exhibiting reciprocal CAs. In 3 out of these pairs, we find evidence for two separate CAs repeatedly affecting the same respective haplotype one after the other, respectively (**Fig. 2B, S2B**). This implies a multi-step process^3,6^, with CAs arising across successive cell divisions, and allowed us to resolve the respective rearrangement history. To exemplify this, **Fig. 2B** depicts reciprocal CAs that based on genomic reconstruction arose in consecutive BFB^19,39^ cycles. In the first cell cycle, sister chromatid fusion followed by bridge-breakage gave rise to two cells carrying an inverted duplication on chromosome 14 as well as a terminal deletion affecting the same homolog (**Fig. 2C**). In the second cell cycle, the cell carrying the terminal deletion underwent a second fusion and bridge-breakage resolving to the same haplotype. Notably, we also find evidence for complex CAs arising during the most recent cell cycle near the second bridge-breakage site (red tick marks in **Fig. 2B**), as discussed further below. Collectively, these CAs gave rise to sister cells harbouring a shared terminal deletion, as well as reciprocal gain-loss events. The reconstructed succession of CAs is supported by three orthogonal data layers^39^– complementary read-depth, reciprocal template strand segregation, and the reciprocal redistribution of haplotype segments. These data highlight how MAGIC detects mitotic errors and how its coupling with Strand-seq enables reconstructing CAs arising across successive cell cycles.

#### Analysis of the de novo CA landscapes of two near-diploid cell lines

With the link between micronucleation and CA formation verified, we next investigated the landscape of CAs arising in conjunction with spontaneous micronucleation, initially focussing on MCF10A (**Fig. 2D**). We find that the most common CA class comprises a simple gain or loss of terminal chromosome segments, representing 21.8% and 19.4% of CAs, respectively. Simple, interstitial CAs appear to be much more infrequent, representing only 4% of the CAs detected from Strand-seq. We also find 29 CAs (23.4%) that appear to have arisen from rearrangements of terminal chromosome sequences over multiple cell cycles (‘terminal multi-step’; see **Fig. S4B**). In 25 single cells with sufficient haplotype support^39^, we find that these CAs consistently arose on the same homolog, in support of multi-step CA formation. Additionally, we find 18 examples of clustered CAs unrelated to terminal multi-step events, likely formed by a complex CA formation process. Leveraging the haplotype resolution of Strand-seq, we confirm that the respective CAs are on the same homolog in line with complex DNA rearrangements, except for one instance where both homologs are affected. These complex CAs include 4 cases of chromothripsis^13,41^ with extensive rearrangements spread across a single homolog (**Fig. S4E**). Lastly, we find that whole-chromosome aneuploidies represent 16.9% of all CAs, with chromosomal losses (*N*=19), interestingly, being much more frequent than gains (*N*=2).

Analysis of the CA landscape in micronucleated RPE-1 cells revealed a similar spectrum of CAs, with terminal chromosomal alterations occurring most frequently. Like for MCF10A, we find that whole chromosome losses (*N*=6) are more common than gains (*N*=1) in RPE-1, a trend that is evident also in normal (control) cells from both cell lines (**Fig. 2E, S4D**). Yet, unlike in micronucleated MCF10A, we observe that complex CAs are essentially absent in RPE-1 cells (**Fig. 2E**, **S4C**). These data underscore the diversity of CA classes emerging in spontaneously micronuclei cells, with evidence for *de novo* complex CAs essentially restricted to micronucleated MCF10A cells in our experiments.

#### Chromosomal distribution and genomic contexts associated with de novo Cas

A number of genomic properties such as replication timing, common fragile sites, and G-quadruplex (G4)-forming sequences have previously been associated with the density of SVs in primary cancer genomes^16,42,43^. We explored the extent to which MAGIC allows evaluating these properties for *de novo* CAs – before Darwinian selection can act – to potentially elucidate genomic contexts susceptible to CA formation. In analysing the genome-wide distribution of CAs from spontaneously micronucleated cells, we find no enrichment for fragile sites. Yet, we observe overrepresentation of sequences forming G4-quadruplex for both CA breakpoints and SCEs, the latter of which mark sites of DNA double-strand breakage (**Fig. S5A**; *adj. P*=0.012, permutation test). We also observe enrichment for CA breakpoints in both early and late-replicating regions, accompanied by a similar trend for SCEs (**Fig. S5A;** *adj. P*<0.05).

We next examined the *de novo* CA density across each cell line’s chromosome set in spontaneously micronucleated cells. While we do not find evidence for recurrent breakpoints, we observe an uneven density of CAs across the MCF10A karyotype. Under the assumption that each homolog acquires CAs with equal probability, we find an enrichment of CAs on chromosome 19 (*adj. P*<0.05, binomial test; **Fig. 2F**). Since most CAs are terminal to a chromosome arm and thus involve telomeres, we conducted Oxford Nanopore Technologies (ONT) long-read sequencing on an MCF10A-derived clone (clone 7; **Methods**), with ONT reads enabling telomere length inference^44^. Analysing the ONT reads, we observe a significant inverse correlation between the density of CAs per chromosome arm and telomere length estimates from Telogator^44^ (Pearson’s *R*=-0.39; *P*=0.0073), with arms most often impacted by CAs exhibiting particularly short estimated telomere lengths (**Fig. S5B**). This suggests that shortened telomeres^4,45–47^ can foster mitotic errors resulting in CA formation in MCF10A.

By comparison, in RPE-1 we find that chromosomes 2, 6, 10, and X each exhibit elevated *de novo* CAs (**Fig. 2G**, *adj. P*<0.05). Overall, we note a bias towards CAs originating from larger chromosomes in RPE-1 (*P*<0.05), a trend not seen in MCF10A (**Fig. S5C**). This could result from differences in the tendency of large versus small chromosomes^48–50^ to be included in micronuclei between both cell lines. In RPE-1, the highest CA counts are seen for chromosomes 10 and X, which engage in an unbalanced der(X) t(X;10) translocation in this cell line^39,51^ (**Fig. 2G**). These data suggest that the der(X) t(X;10) derivative chromosome in RPE-1 cells might be particularly susceptible to inclusion in micronuclei, which could promote its genetic instability.

#### Distinct chromosomal distribution patterns in clonally propagated versus de novo Cas

While our data so far imply that non-transformed cells undergo continuous karyotype changes, MCF10A and RPE-1 cell lines maintain a relatively stable karyotype in culture. To assess the potential of CAs to clonally propagate, we employed MAGIC to isolate micronucleated cells, testing their ability to form new clones. We find a reduced success rate in generating clones from micronucleated cells compared to normal cells, in both MCF10A and RPE-1 (1.3 and 2.5-fold respectively, *P*<0.001, Fisher’s exact test; **Fig. S7D**). To explore the ability for CAs to persist clonally, we expanded 18 and 7 clones from MCF10A micronucleated and control cells, respectively, followed by analysis through bulk-cell low-pass WGS (**Methods**). We find 13 clonally propagated CAs in the clones expanded from micronucleated cells (**Supplementary Table 1**), with 9 out of 18 seeded cultures containing at least one clonal CA – which appear to represent both simple (*N*=12) and complex (*N*=1) events, based on the chromosomal clustering of CAs inferred by read depth analysis. By comparison, 3 clones grown from the controls each contain one simple CA. By pursuing ONT sequencing of one clone deriving from a micronucleated cell (clone 7), we verify complex CA formation, by identifying a chromothripsis event that followed isochromosome formation (**Fig. S7G, S7H**). These data show that MAGIC can be employed to efficiently select cells subject to CAs for phenotypic analysis and clonal expansion.

In comparing clonally propagated CAs with the *de novo* CAs spectrum in MCF10A, we observe no CAs affecting chromosome 19 after clonal growth. Additionally, we note a prevalence of losses on the 7q-arm, involving simple and complex CAs and comprising 5/13 (38.5%) of all clonally propagated CAs—a 4-fold enrichment when compared to *de novo* CA spectrum from the same cell line (*P*=0.0043; Bonferroni corrected Chi-square test; **Fig. S7F, S7E**). While 7q-losses are common in breast cancer^8,52^ (see **Supplemental Note**), these CAs may either be subject to positive selection or could have persisted as selectively neutral events during clonal expansion. Overall, these observations are consistent with differences in the genomic spectra of clonally propagated and *de novo* CAs, likely influenced by selective forces^3^ acting during clonal growth.

### Prevalence of dicentric-mediated CAs and chromothripsis in spontaneously micronucleated cells

#### Pivotal role of dicentric chromosomes in spontaneous CA formation

Capitalising on the utility of Strand-seq data to discern distinct CA classes^39,40^, we next systematically inferred *de novo* CA processes, combining the single-cell genomic data generated for both cell lines. We first assessed terminal *de novo* CAs, which represent 64.6% and 75.5% of all CAs seen for MCF10A and RPE-1, respectively, and which are equally distributed between gains and losses. Out of the 49 terminal gains observed in both cell lines involving either parts of or an entire chromosomal arm, 42 (85.7%) show a configuration where the segment gained at the terminus has a strand-state opposite to its source homolog, consistent with a terminal inverted duplication^39^ (**Fig. S6A**). This karyotypic pattern could arise from an initial BFB cycle stage, yet may alternatively reflect acentric fragments entering mitosis unrepaired and undergoing asymmetric segregation (**Fig. S6C**).

To distinguish between both scenarios, we considered that reciprocal gain-loss patterns would emerge in sister cells according to both mechanisms. Yet, BFB cycles would be confined to a single homolog^19,39^, and would be prone to show sequentially forming CAs affecting that homolog. When analysing CAs locating to the same chromosome, whereby at least one CA involved its terminus, 86.2% originate from the same homolog, consistent with BFB cycles (**Fig. 2H**; **Fig. S6B**). These data are further bolstered by our analysis of three sister cell pairs harbouring at least one shared and one reciprocal CA affecting the same homolog, in each case verifying multi-step BFB cycles (**Fig. 2B**, **S4B**). Collectively, these data underscore the important role of dicentrics in facilitating BFB-mediated CAs in these cell lines, while also suggesting that additional CA processes—discussed in the subsequent sections—contribute to their spontaneous *de novo* CA landscapes. At least in MCF10A, telomere shortening may commonly trigger spontaneous CAs arising via dicentric chromosomes.

#### Complex CAs arising in conjunction with spontaneous dicentric-mediated de novo rearrangements

We next focused on additional *de novo* CA processes. Our analysis of CA patterns revealed two distinct types of complex CA, in each case implicating chromothripsis. Among all 54 CAs involving a terminal deletion or inverted duplication in spontaneously micronucleated MCF10A cells, we find that 11 (20.3%) exhibit a localised small-to-intermediate scale copy-number oscillation pattern near the internal breakpoint strongly indicative of complex CA formation (**Fig. S6D, S6E**). This CA pattern is further corroborated by its similar occurrence frequency in MCF10A control cells, and can be also seen in RPE-1 cells (**Fig. S6E**). Pooling examples of this pattern across all examined conditions, the Strand-seq data indicate a single oscillation in the majority of cases (11/16, 68.7%), characterised by troughs and crests of similar size, averaging 1.6 Mbp, with the whole pattern spanning from 2 to 7 Mbp (**Fig. S6F, S6G, S6H**). Assuming chromatin bridge breakage as the source of these CAs, the location of these small-scale complex rearrangements corresponds to the point of rupture of the affected homolog, suggesting a link to bridge resolution. The location and oscillating nature of this CA pattern mirror previously reported cases of chromothripsis linked to dicentric breakage, following the disablement of the Rb and p53 pathways in RPE-1 cells^4^. Cytosolic enzymes like the Three Prime Repair Exonuclease 1 (Trex1)^4^ potentially resulted in these spontaneous complex CAs.

#### Whole chromosome pulverisation in spontaneously micronucleated cells

In MCF10A cells, but not in RPE-1, we further detect 4 instances of whole chromosome or arm-level chromothripsis. In each case, all observed rearrangements are confined to a single haplotype and show evidence for random fragmentation along the homolog, in line with established chromothripsis criteria^13,41^ (**Fig. S4E, 2I, S7C**). In 2 instances, we find the corresponding sister cell in our data via single-cell genomic analysis (in one case this included one sister cell not initially passing quality control). Analysing both sister cell pairs, we observe anti-correlated read counts indicating reciprocal segregation of pulverised fragments (**Fig. 2I, S7A, S7B, S7C**), thus independently verifying the occurrence of chromothripsis. These spontaneous CA patterns mirror reports of chromothripsis linked to micronucleus entrapment, as previously demonstrated in *TP53*-depleted RPE-1 cells after monastrol washout^5^. Considering both chromothripsis patterns – localised and chromosome-wide – chromothripsis appears to have contributed to up to 13% of the alterations seen within the CA landscape of spontaneously micronucleated MCF10A cells.

### Disentangling the effects of p53 status on CA formation processes

#### Effect of TP53 status on de novo CA formation

Disruption of *TP53*, causing loss of the p53 tumour suppressor, is the most common driver mutation in cancer, and associated with a range of genomic instability patterns including chromothripsis^14,53–55^. However, the role of *TP53* deficiency in increasing the baseline CA formation rate or altering the *de novo* CA spectrum has remained challenging to determine. To explore these questions, we utilised previously developed *TP53-/-* MCF10A and *TP53-/-* RPE-1 cells^56^ (**Fig. S8A, Methods**). Using confocal microscopy, we find a general increase in nuclear atypias for both *TP53-/-* cell lines compared to their unmutated (WT) counterparts (**Fig. S8B, S8C**). Exemplifying this, about a third of *TP53-/-* MCF10A cells exhibit micronuclei (**Fig. 3A**), an increase that appears to be mainly due to a higher frequency of anaphase bridges, reaching 36.8% of all cell divisions (**Fig. 3B**). Moreover, the probability of anaphase errors to result in a micronucleated daughter is increased to 73.3% and 69.7% for anaphase lagging chromosomes and chromatin bridges, respectively (**Fig. S8D**). Similar to their WT counterparts, micronucleated *TP53-/-* cells are prone to generate a micronucleated daughter cell (**Fig. S8E, S8F**). Yet, unlike WT cells, the duration of the cell cycle is not prolonged, and these *TP53-/-* cells do not effectively enter cell cycle arrest (**Fig. S8G, S8H**), in accordance with established roles of p53.^53^ These observations hint at an elevated rate of spontaneous CAs in *TP53-/-* cells, with the absence of efficient cell cycle arrest upon micronucleation potentially promoting CA formation.

**Figure 3:**
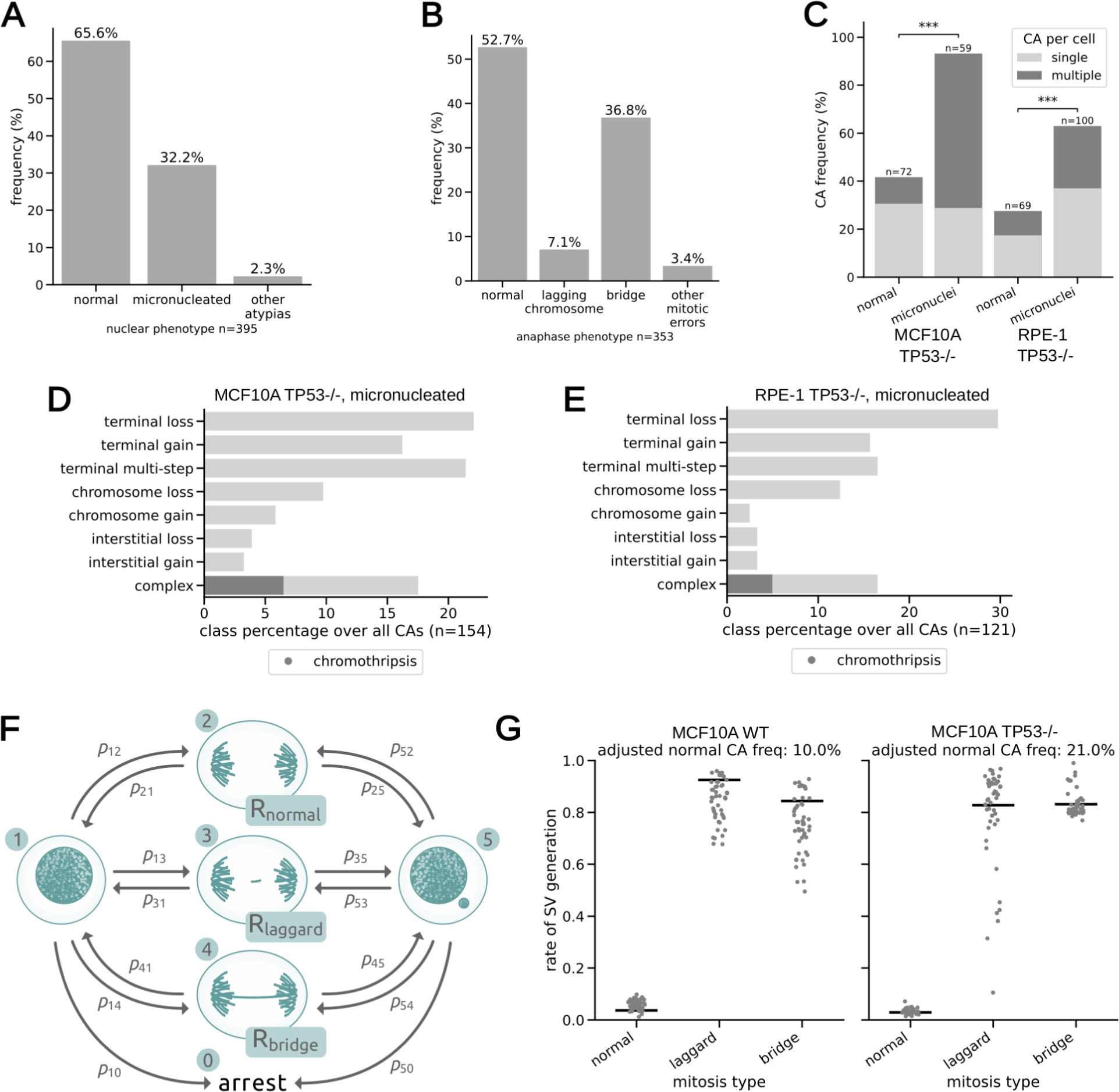
Effect of TP53 disruption on de novo CA formation, and modelling basal CA rates. (A,B) Frequency of nuclear (A) and anaphase (B) phenotypes in MCF10A TP53-/-cells from long-term live-cell imaging. (C) Observed CA frequency by cell. (D,E) Breakdown of CA classes in micronucleated MCF10A TP53-/- (D) and RPE-1 TP53-/- (E) cells, class percentage over all CAs. (F) Agent-based model for estimating the de novo CA rate. Scheme representing available states (numbers in teal circles) and available transitions (arrows) between states, with p_ij_ being the empirically measured transition probabilities. Parameters are estimated for three mitosis types with R representing estimated CA rates. 1, normal cell; 2, normal mitosis; 3, laggard mitosis; 4, bridge mitosis; 5, micronucleated cell. (G) Mitosis type-specific CA rate estimations in MCF10 WT and TP53-/- (KO) models. Each data point represents the estimated value from an optimisation run. Black lines represent the average weighted by the residual error of each optimised simulation.

To investigate the effect of *TP53* disruption on *de novo* CAs, we subjected both *TP53-/-* cell lines to MAGIC followed by Strand-seq. Our analysis of the 300 single-cell genomes generated demonstrates a marked increase in *de novo* CAs (**Fig. 3C**), with *TP53-/-* micronucleated cells showing considerably higher CA numbers compared to WT micronucleated cells in both cell lines (compare with **Fig. 3C, S9A**; *P*<2.65e-10 for MCF10A; *P*<9.36e-6 for RPE-1). We conducted an analysis of CA classes arising in both *TP53-/-* cell lines compared to their WT counterparts (**Fig 3D, 3E, S9B, S9C**). While we find the CA spectra to be very similar between *TP53-/-* and WT (**Fig. S9D**), a notable exception is the marked increase in complex CAs emerging in micronucleated *TP53-/-* RPE-1 cells compared to WT (from 2.7% to 16.5%; *P*<0.05 Fisher’s exact test; compare **Fig. 2E** and **Fig. 3E**), including chromothripsis events. This indicates that while *TP53-/-* cells do not generally show an altered *de novo* CA spectrum, complex CA formation is influenced by *TP53* deficiency under certain conditions.

#### De novo CA rate estimates in normal and p53-deficient MCF10A cells

Determining the baseline rate of somatic CAs formation has remained challenging due to previous technological limitations^3^. Capitalising on the large-scale data generated in our study, we devised a statistical agent-based model (**Methods**) where simulated cells transition between nuclear atypias and normal mitoses based on probabilities derived from live-cell long-term imaging (**Fig. 3F, 1B**). This model allows simulating the rate of *de novo* CA associated with three mitosis types: normal, with lagging chromosomes (‘laggard’), and with chromatin bridges (‘bridge’). We focused on MCF10A, because our data show CAs actively arise in both *TP53-/-* and WT contexts in this cell line. We simulated a population of cells, allowing each cell to progress through mitosis and, upon division, acquire a new nuclear phenotype and CAs according to the model’s transition probabilities. At the end of the simulation, we measured the error between the inferred CA rate and the empirically observed CA frequency in normal and micronucleated cells. An optimization algorithm minimising this error allows estimation of the basal CA rate. Encouragingly, despite not being explicitly programmed into the model, the frequency of micronuclei stabilised at 5.0% and 38.3% for WT and *TP53*-/-cells respectively, closely mirroring our empirical data (**Fig. S10A, S10B**).

Notably, the CA events detected in a cell population reflect a combination of newly arisen (*de novo*) CAs and those inherited from earlier cell divisions. To account for this, we evaluated the goodness-of-fit by adjusting empirically observed CA counts across a range of assumed *de novo* CA proportions (**Methods**). After applying this adjustment, the optimal fit for normal nuclei corresponds to a 10% *de novo* CA proportion in WT cells and 21% in *TP53-/-* cells. With the same adjustment parameter applied to micronucleated cells, 47% of CAs in WT cells and 72% in *TP53-/-* cells represent *de novo* events (**Fig. S10C**). Using these adjusted parameters, we first estimated the mitosis type-specific CA rate, which is 3.7% per cell division for WT cells undergoing a normal mitosis. By comparison, mitosis type-specific CA rates are 92.5 and 84.4% per cell division for laggard and bridge mitoses in WT-cells, respectively (**Fig. 3G, S10D, S10E**). In *TP53-/-* cells, mitosis type-specific CA rate estimates are very similar – 2.9% for normal, 82.8% for laggard and 83.2% for bridge mitosis per cell division – suggesting that the underlying processes by which CAs form via nuclear atypias are not affected by *TP53* deficiency. Finally, by considering the relative contribution of mitosis types, we estimate the basal CA rate for MCF10A, which is 13.3 % in WT cells per cell division, and approximately doubles to 30.4% in *TP53*-/-cells. This increase appears mainly driven by the higher proportion of cells bearing nuclear atypias – from 11.3% in WT to 43.9% in *TP53-/-* cells – with a particularly pronounced contribution of chromatin bridges (**Fig. 3B**), consistent with dicentrics being an important trigger for sporadic CA formation.

### Chromosomal location determinants for CA formation

#### Modelling CA formation determinants with targeted DSBs results in diverse SV spectra

CA drivers surpass base substitution drivers in number, yet the role of DSB positioning along chromosomes in triggering CAs is under-explored^3,16,37^. Based on our data indicating dicentrics are an important trigger for CAs, we revisited our data from the X chromosomal *HPRT1* locus. We find that the majority (53.5%; 23/43) of the segmental CAs affecting chromosome X display alterations of centrally-oriented segments relative to the cut site, indicative for BFB cycles appearing to lead to repeated DSBs triggering *de novo* CAs. Based on these data, we reasoned that two factors are likely to determine CA formation during micronucleus entrapment – sister-chromatid fusions, and the fate of chromosome fragments during mitosis. Thereby, the size and nature of the fragments generated, and whether they are dicentric or acentric (**Fig. 2H, S6C**), are likely to influence their fate. This implies that the positioning of DSBs along chromosomes may play an important role in CA formation. We reasoned that the capabilities of MAGIC would allow exploring this question systematically.

We thus devised a MAGIC experiment generating DSBs on the q-arms of chromosomes 2 and 7 using CRISPR/Cas9. Each chromosome was targeted at a specific sub-centromeric, sub-telomeric, and central site on the q-arm (**Fig. 4A**). We selected chromosome 2q due to its low average repeat content facilitating gRNA design, and 7q due to the enrichment for clonally propagated CAs we observe for this arm. After subjecting MCF10A cells to targeted DSBs followed by micronucleated cell selection, we sequenced 361 single-cell genomes from these MAGIC experiments (**Supplementary Table 2**). While we observe different CA induction efficiencies for each DSB target region (ranging from 20 to 60%), in each case the majority of CAs originate from the gRNA-directed DSB sites (**Fig. 4C**), once again verifying that MAGIC effectively captures CAs triggered through targeted DSBs.

**Figure 4:**
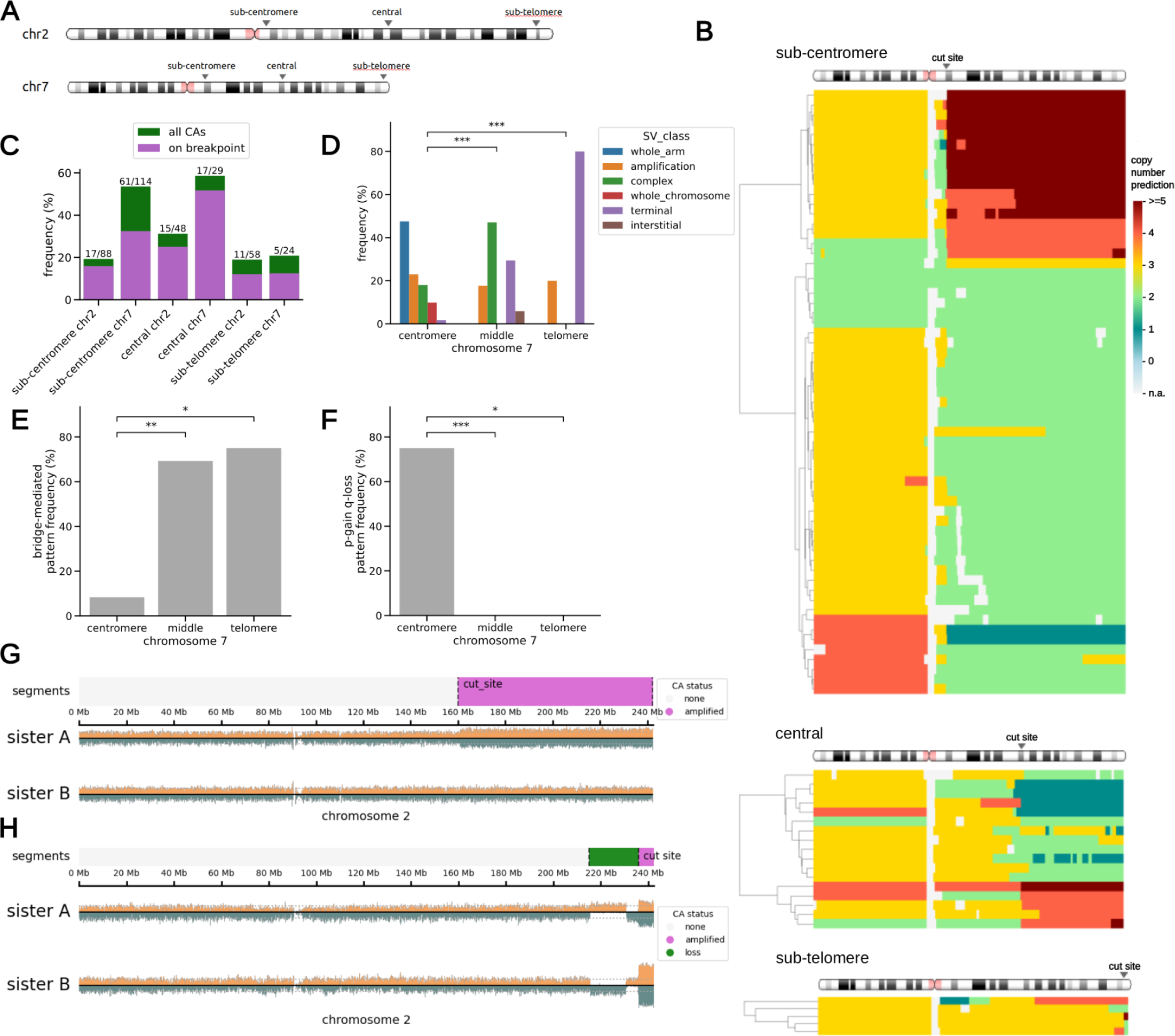
CA landscape following targeted DSB induction along chromosome arms. (A) Scheme displaying the locations of targeted DSBs (B) Overview of CAs for the chromosome 7 q-arm, expressed as copy-number states. Sub-centromere, central, and sub-telomere target loci are depicted from top to bottom. (As a note, the consensus copy number for chromosome 7 in MCF10A H2B-Dendra2 cells is three - trisomy). (C) CA frequency per cell across the different chromosomal cut sites. (D) Frequency of CA types for different cut sites on the chromosome 7 q-arm. (E,F) Enrichment for bridge-mediated CAs (E) and coupled p-arm-gain/q-arm-loss indicative for isochromosome formation (F). Only terminal and complex CAs were considered in this analysis. (G,H) Sister cells displaying one-sided (G) and asymmetric (H) inheritance of the acentric fragment. The sister cell examples shown here are from the chromosome 2 central cut and HPRT1 datasets, respectively.

We next analysed the *de novo* CA spectra separately by DSB site. We observe a wide diversity of CA classes (**Fig. 4B, 4D, S11F**), including cases of chromothripsis as previously reported after targeted DSB induction^37^. Yet, the relative composition of CA classes observed markedly varies by the cut location, with trends largely consistent across 7q and 2q (**Fig. 4B, 4D, S11A, S11B**). In particular, we find terminal CAs affecting the q-arm – seen with 29.4% and 80.0% for 7q, for example – when specifically targeting the central and sub-telomeric site, respectively, whereas whole-arm CAs arise exclusively from sub-centromeric DSBs. An in-depth analysis of CA outcomes reveals that these patterns reflect not just the size of the generated fragment, but imply the activity of distinct CA processes. Particularly, when focusing on those CAs initially annotated as either terminal or complex, we find several examples of terminal deletions with an inverted duplication centrally-located relative to the DSB (**Fig. 4B, S11C, S11A**), consistent with dicentric-mediated BFB cycles. These bridge-related CAs are >8-fold enriched in central and sub-telomeric cuts, with significant enrichment compared to sub-centromeric cuts replicating between 7q and 2q (**Fig. 4E, S11D**). Since bridge-related CAs can trigger genetic instability over successive mitoses, this suggests that the activity of certain CA processes is influenced by the initiating DSB location.

Furthermore, when targeting the sub-centromeres, we observe a notable frequency – 10.5% and 9.1% for 7q and 2q, respectively – of whole-arm CAs affecting both the targeted long arm in *cis* and the untargeted short arm in *trans* (**Fig 4B, S11A**). A relevant subset of these shares a distinctive CA pattern characterised by a p-arm gain in inverted orientation coupled with q-arm loss, indicative of isochromosome formation (**Fig. S11G**). By comparison, neither central nor sub-telomeric cuts result in isochromosomes (**Fig. 4F, S11E**), implying that sub-centromeric DSBs are capable of inducing isochromosome formation—resulting in a derivative chromosome structure recurrent in different cancer types such as medulloblastoma and lung cancer^8,57^. Taken together, these data provide strong evidence that different DSB sites can trigger distinct *de novo* CA processes, depending on the DSB location along the chromosome.

#### Amplification of large acentric fragments appears to mediate their co-segregation

Across cut sites, we also observe several cases of amplification of the acentric fragment generated from DSB induction. This pattern accounts for 17.6 and 40.0% of all CAs, depending on the site, with the amplifications reaching up to 142 Mb in length (**Fig. S12A, S12B, S12C**). Unexpectedly, Strand-seq analysis revealed simultaneous gain of both Watson (W) and Crick (C) templates^38,39^ for the region encompassing the acentric fragment, with W/C ratios converging at 1:1 (**Fig. S12D**): Among 18 acentric gains with a copy-number increment of 2, the W/C ratio remains 1:1 in all 18 cases, with C/C and W/W configurations missing (*P*<7.63e-06, binomial test; **Supplementary Methods**). This peculiar strand pattern implies that the acquired acentric segments are integrated into the same derivative chromosome, promoting their co-segregation in multiples of two, with the segments being arranged in an inverted orientation (**Fig. S12E**). This inference is further corroborated by sister cell pair analysis, demonstrating the joint segregation of gains in multiples of two: in one instance, we observe one-sided segregation of the amplified fragment, including both the W and the C strand (**Fig. 4G**); in another, we find asymmetric inheritance of the acentric amplification among sisters, again involving W and C template co-inheritance (**Fig. 4H**). In summary, coupling MAGIC with CRISPR/Cas9 provides evidence for a CA process promoting the co-segregation of amplified acentric DNA segments.

#### Fluorescent in situ hybridization reveals isodicentric chromosomes as well as acentric fragment co-segregation through isoacentric formation

To further investigate these reconstructed CA patterns, we conducted an additional round of targeted DSB experiments, this time coupled with karyotyping by fluorescent *in situ* hybridization (FISH). We utilised a two probe strategy labelling the sub-centromeric regions of 7p and 7q, respectively. The gRNA cut site, thereby, is located within our designed q-arm FISH probe, facilitating the analysis of CA outcomes since rearrangements at the cut site will effectively ‘split’ the probe signal (**Fig. 5A**). Following centromeric cuts, we find that 28% of metaphases display an abnormal sub-centromeric probe signal indicative for CAs, an outcome never observed for sub-telomeric cuts (**Fig. S13A**).

**Figure 5:**
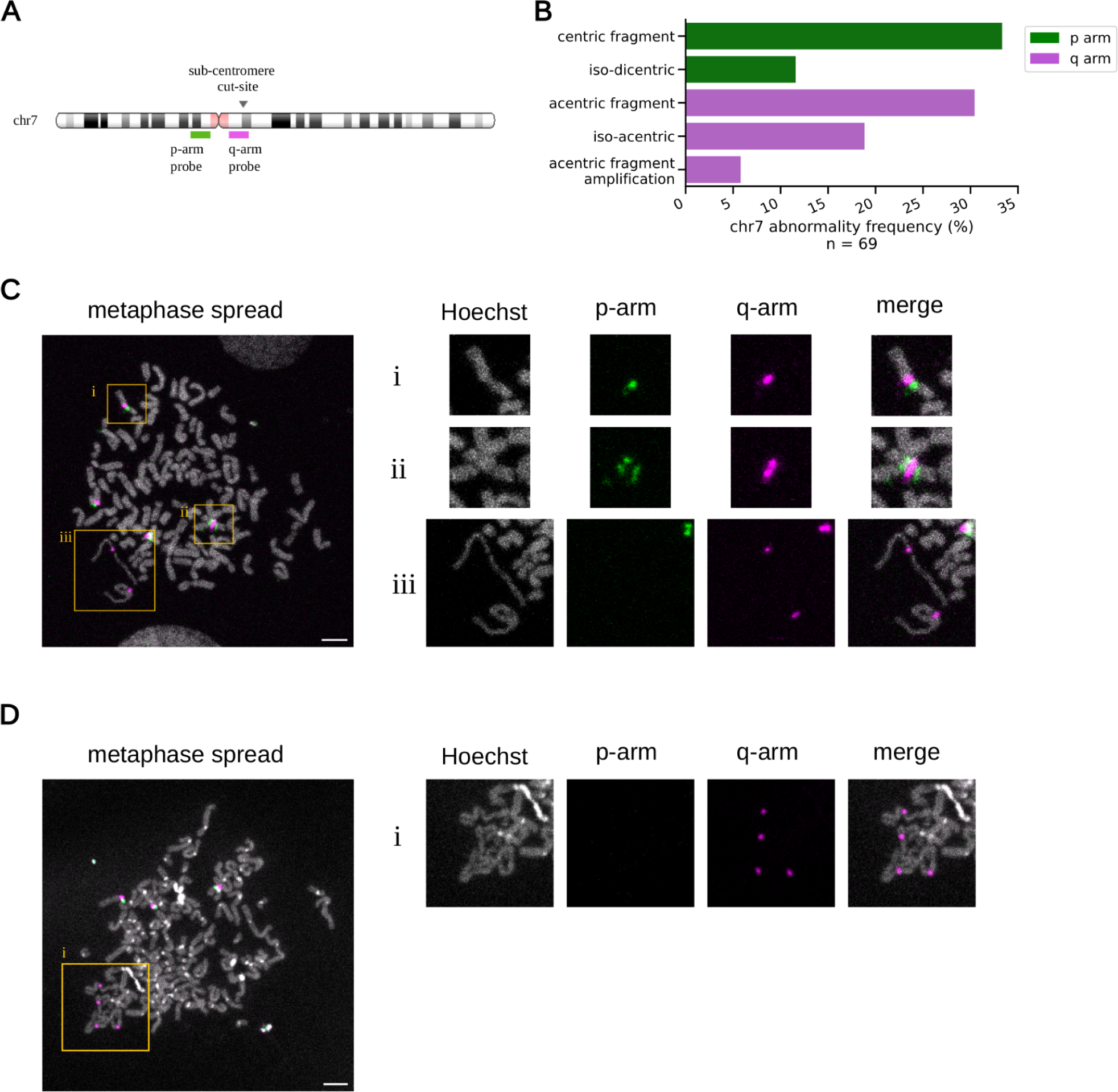
Isodicentric and isoacentric derivative chromosome generation through targeted DSBs. (A) Scheme depicting hybridisation sites for sub-centromeric p-arm and q-arm FISH probes. The targeted DSB site, ‘splitting’ one of the FISH-probe locations, is indicated by a black arrow. (B) FISH-based quantification of chromosome 7 abnormalities displaying an abnormal probe pattern in metaphase spreads (C) Metaphase spread example revealing different derivative chromosomes. Left: regions of interest (ROIs) are marked by yellow squares. Right: magnified ROIs, with breakdown per fluorescent channel. ROI i: normal chromosome 7. ROI ii: isodicentric chromosome with a q-arm signal surrounded by two p-arm signals, indicating the presence of two adjacent centromeres and duplication of 7q. ROI iii: isoacentric derivative chromosomes with signs of premature chromatin condensation. (D) Metaphase spread example with amplified isoacentrics. ROI i: visibly clustering, amplified isacentrics, with several interspersed q-arm signals clearly visible.

An in-depth analysis of each metaphase spread, notably, demonstrates patterns of CA verifying those genomically reconstructed using Strand-seq: Following sub-centromeric cutting, we observe a loss of the long arm at the DSB site in 33.3% of all spreads with an abnormal chromosome 7 (centric fragment, **Fig. 5B**). Additionally, we find that isochromosomes—more specifically isodicentrics—account for 11.6% of all abnormalities, as visualised by two sub-centromeric p-arm signals surrounding a single sub-centromeric q-arm signal (**Fig. 5B, 5C**). These data validate isochromosome formation following targeted DSB generation, resulting in a derivative chromosome that despite comprising two centromeres is likely to represent a genetically stable structure.

Our FISH experiments further highlight multiple abnormalities affecting acentric fragments. Acentrics appear as an isolated chromosome fragment, with a q-arm signal at one extremity, in 30.4% of spreads with an abnormal chromosome 7 (**Fig. 5B, S13B**). Notably, in 18.8% of cases, we detect 7q acentric fragments that have doubled in size, bearing a sub-centromeric q-arm probe signal located at the middle (**Fig. 5B, 5C, S13C**). These derivative chromosomes represent isoacentrics, showing a pair of mirrored acentric arms that intersect at the initial cut site without the presence of a centromere. This pattern harbours the acentric segment copies in inverted orientation, again demonstrating validation of our genomic reconstructions. Notably, these isoacentrics appear thin and elongated, suggesting that they could be subject to abnormal chromatin condensation (**Fig. 5C**). Such morphology has been previously associated with premature chromatin condensation^58,59^, and could reflect under-replication due to micronucleus entrapment^59^. Interestingly, in 5.8% of metaphase spreads with an abnormal sub-centromeric probe signal, we observe further amplified isoacentrics, which present as clusters of condensed DNA with interspersed sub-centromeric q-arm signals (**Fig. 5B, 5D**). It is intriguing to speculate that these condensed DNA structures might further promote the co-segregation of amplified acentric genetic material in multiples of two. These findings highlight how targeted DSBs can help to systematically elucidate CA formation processes using the MAGIC platform. They further underscore the critical role of sub-centromeric regions in somatic CA formation processes.

## Discussion

Following Boveri’s early 1900s research linking abnormal mitoses to cancer, scientists have long sought to unravel the processes leading to somatic CAs, with genetic instabilities resulting in chromosomal alterations constituting an enabling hallmark of cancer^60^. Yet, the rate at which somatic CAs arise in cells has remained poorly understood^3^. By integrating automated imaging and genomics with the autonomous MAGIC platform, we have systematically captured CA formation in near-diploid human cells, providing insights into *do novo* CA processes and rates.

### Role of dicentric-mediated CAs, prevalence of chromosome losses, and cell-line specific differences in *de novo* CA spectra

Unlike prior studies that induced nuclear atypias to enable studies of CA formation processes, MAGIC allows investigating CAs arising in conjunction with spontaneous nuclear atypia formation. This is likely to provide a more accurate representation of the variety of CAs arising in human cells. Our analyses highlight dicentric chromosomes as prevalent triggers^6,61^ of spontaneous CAs. The ability of dicentrics to trigger CAs clustering on a single homolog over multiple cell cycles underscores their importance in mediating somatic karyotype diversification.

Our data further show a bias in *de novo* aneuploidy generation towards chromosome losses that arise more often than gains. These data are bolstered by the KaryoCreate system, which utilises mutant kinetochore proteins to create aneuploid cells, and reportedly shows a 50% higher efficiency for generating chromosome losses vs. gains after clonal propagation^62^. Our data are also paralleled by the prevalence of chromosome losses over gains in cancer genomes, initially observed with microarrays^63^. Notably, our analysis of 2,600 WGS pan-cancer genomes^8^ (**Supplementary Notes)** reveals that losses constitute most (81.5%) aneuploidies in cancer even when excluding tumours subject to whole genome duplication^64^ (**Fig. S14A**). Together with our observation that *de novo* structural aberrations in MCF10A resemble patterns of CA observed in primary cancers (**Supplementary Notes**) – particularly CAs that appear to emerge within a largely disomic context^8,22^ (**Fig. S15**) – these data suggest that CAs arising in micronucleated cells can markedly shape cancer genomic landscapes.

The mechanistic basis for the biassed formation of chromosome losses over gains is not immediately clear. Prior reports highlighted roles of proteotoxic stress associated with somatic trisomy^65,66^, which could result in the preferential selective removal of chromosomal gains. Yet, our data imply that a bias towards chromosome losses is established already during CA formation, a phase where proteotoxic stress may be less likely to represent a crucial factor. Factors contributing to chromosomal loss bias could include defective replication and breakage of chromosomes within the micronucleus^32^. Additionally, segregation of dicentrics to a single daughter cell^67^ could have resulted in monosomy in one daughter and introduced a derivative chromosome susceptible to further CAs in the other. Finally, the elimination of micronuclei through degradation or extrusion from cells^68,69^ may contribute to the preferential formation of spontaneous chromosome losses determined by the MAGIC platform.

Our comparative analysis of MCF10A and RPE-1 cells reveal certain differences with respect to *de novo* CA formation. MCF10A cells exhibit *de novo* CAs across their entire karyotype, with a notable tendency for complex CA formation. This, notably, includes documentation of *de novo* chromothripsis in cells with a normal *TP53* gene^70^, and thus indicates that chromothripsis can occur *de novo* at an appreciable rate even when *TP53* is intact. Initiating immortalisation events in MCF10A,^71^ including genetic gain of the *c-Myc* gene as well as the combined loss of *CDKN2A* and *CDKN2B*, may have enabled frequent sporadic *de novo* chromothripsis in a p53 competent context. By comparison, RPE-1 exhibits a more stable karyotype, few complex CAs, no chromothripsis, and a preference for *de novo* CAs affecting large chromosomes. While absent in WT RPE-1 cells, RPE-1 *TP53^-/-^* cells exhibit complex CA formation including chromothripsis, mirroring the previously reported link between chromothripsis and mutant *TP53* in tumours such as medulloblastoma^14^. Collectively, these data underscore the context-specific nature of *de novo* CAs, with potential roles of intrinsic cell type characteristics such as cell cycle regulation and DNA repair pathway activity, in CA formation. The tendency of larger chromosomes to form micronuclei more frequently in RPE-1 cells^48–50^, an apparently genetically unstable der(X) t(X;10), and telomere length variations are likely to explain additional cell line-specific differences in *de novo* CA spectra observed.

### Influence of the chromosomal position of the initiating DSB on CA formation

By integrating CRISPR/Cas9 engineering with MAGIC, we have delineated the potential for DSB sites to trigger CAs, with distinct CA outcomes emerging depending on where the initiating DSB occurs. This includes cases in which the resulting derivative chromosomes, particularly isochromosomes, can be stably inherited over successive cell cycles, and those where the derivative chromosome appears unstable resulting in further CAs. Our experiments demonstrate that isochromosomes form by a process initiated from sub-centromeric DSBs, which is followed by sister chromatid fusions likely taking place in the micronucleus. This process locks the mirror-arms of the resulting isodicentric chromosome in an inverted orientation. Prior studies had suggested the possibility of isochromosome formation through such a process, coined U-type exchange^72^, yet, to our knowledge definitive experimental evidence for this process was previously lacking.

We note that our data imply that the inter-centromeric distance, defined as twice the distance from the DSB site to the centromere in a dicentric, plays an important role in CA formation (**Fig. 5C, S13D**). If the inter-centromeric distance surpasses a certain threshold, each centromere can attach to a kinetochore, forming chromatin bridges triggering further *de novo* CAs. Conversely, if the inter-centromeric distance is shorter than the threshold, the two centromeres would be handled as a single unit, resulting in a single kinetochore attachment and thus enabling stable segregation during mitosis. Indeed, a prior study reported that isodicentrics are remarkably stable, even with large intercentromeric distances as estimated by FISH^73^. Consistent with this, our analysis of primary cancer genomes verifies that isodicentrics arise recurrently in tumours: From 31 up to 55% of isochromosomes present two centromeres in the Cancer Genome Atlas (TCGA) and pan-cancer analysis of whole genomes (PCAWG) WGS resources^8^, respectively, whereby the inter-centromeric distance often surpasses 20 Mb (**Supplemental Note, Fig. S14B, S14C**). Collectively, our findings suggest a crucial role of DSB location in CA formation. They also pave the way for future studies to comprehensively characterise CA processes through targeted DSB generation and MAGIC.

### Isoacentrics facilitate the co-segregation of segmental amplifications: potential implications for DNA rearrangement processes in cancer

Our targeted DSB experiments further reveal the asymmetric segregation of acentric segments amplified in inverted orientation, forming isoacentrics. Among spontaneously micronucleated cells, we observe patterns of template strand co-segregation indicating isoacentric formation in up to 3% of spontaneous micronuclei in both MCF10A and RPE-1 cells (**Fig. S12F, S12G**). By comparison, targeted DSBs result in a marked (∼10-fold) increase in isoacentrics (**Fig. S12H**). This implies that unrepaired DSBs arising within the interior of chromosomes are not the most common trigger of spontaneous CAs in these cell lines. Telomere fusions due to shortened telomeres typically do not generate acentric fragments, and thus trigger CAs through alternative rearrangement processes.

Isoacentrics could result from the fusion of an acentric fragment with its sister chromatid or from an aberrant replicative process, potentially mediating the rapid amplification of DNA segments. Our FISH based metaphase spread analysis shows a shift of the probe signal marking the targeted DSB site to the middle of an isoacentric, at which point this derivative chromosome doubles in size. Therefore, the frequent inheritance of gained chromosomal segments in multiples of two as recently documented through *in vitro* evolution screens^74^ could be explained, at least in part, by isoacentric formation.

We emphasise that the inverted duplication pattern of isoacentrics determined in this study implies that fold-back inversions^75^ could mark a common CA process other than the BFB cycle, thus impacting the interpretation of cancer genomes^8,18,76^. If coupled with other CA processes, isoacentrics may facilitate rapid oncogene amplification and extrachromosomal circular DNA formation, or serve as a precursor for complex CAs. Chromosomal clustering during mitosis, potentially mediated by the glue-like properties of the Ki-67 protein^77^, might enable the asymmetric segregation of intact isoacentrics following amplification of this derivative chromosome structure. Furthermore, the tethering of DNA fragments derived from isoacentrics, potentially mediated by CIP2A and TOPBP1 complexes^25,26^, could contribute to complex CA formation processes. It is interesting to speculate that isoacentrics, if concurrent with chromothripsis, may explain prior reports of chromothripsis with multiple copy-number states^13,14^. Additionally, isoacentrics could potentially contribute source material for chromoanasynthesis—a process characterised by aberrant DNA replication with clustered deletions, duplications, and triplications, often leading to three or more copy-number states^78^.

### Methodological advances, limitations, and implications for future studies

MAGIC builds upon approaches coupling the imaging of visual phenotypes with precise optical tagging^79–81^, by utilising an autonomously operated, ML-assisted adaptive feedback loop designed to systematically investigate *de novo* CA formation. The complementary photolabeling dyes we show to be compatible with MAGIC should in the future facilitate application to a wide spectrum of biosamples. MAGIC enables the analysis and scoring of up to 80,000 cells per experiment to allow isolating cells with rare morphologies at scale. Our study thus addresses an important conceptual gap, overcoming prior limitations that hindered the comprehensive investigation of sporadic *de novo* CAs. We chose Strand-seq for single-cell genome sequencing, to enable karyotypic reconstruction resolved by homolog^39,40^. Our study leveraged MAGIC to isolate 2,036 single cells, and performed single-cell genomic sequencing of 1,330 cells using Strand-seq. This yields an unprecedented data resource for exploring *de novo* CA formation, utilising two cell lines extensively employed in research on chromosomal instability processes. We caution that Strand-seq, due to the intermediate sequence coverage generated, is limited to detecting CAs >200 kb. While this size range encompasses >80% of the SV drivers documented in the PCAWG study^3,8^, certain classes of SV such as <200kb-sized deletions and insertions, as well as CAs arising independently of nuclear atypias, are currently not captured – highlighting an important area for future methods development.

The computational workflows steering the autonomous MAGIC platform, which our study releases openly to the academic community, add to its versatility by facilitating the optimisation of resolution, experimental time, and yield for addressing a wide diversity of research questions in relation to chromosomal instability procsses. Potential future applications of MAGIC include the systematic characterization of mutational signatures generated from *de novo* CAs before Darwinian selection occurs, analogous to prior studies that characterised mutational signatures for sporadically arising base substitutions^20,21^. This could enhance our understanding of the aetiology of structural rearrangements. Future research might also involve the integration of MAGIC with genomic sequencing methods beyond Strand-seq, including single-cell multi-omic techniques, as well as the identification of elevated *de novo* CA formation in clinical samples. This could help identify high-risk precancerous lesions and might improve patient outcomes through timely interventions in the future.

In conclusion, MAGIC reflects an important technological advancement, which we harness to investigate the origins of sporadic CA formation in non-transformed cell lines. Our findings not only highlight the role of dicentric chromosomes as a major threat to genomic integrity and the influence of DSB localization on CA outcomes, but also reveal the basal rate at which CAs emerge, and how this is influenced by *TP53* deficiency. These insights contribute to a deeper understanding of the fundamental processes driving CAs and lay the groundwork for future research aimed at elucidating early stages of tumorigenesis driven through somatic karyotype evolution.

## Acknowledgements

We thank David Pellman and Cheng-Zhong Zhang for providing valuable comments on an advanced version of our manuscript. *TP53-/-* MCF10A cells are a kind gift from Claudia Scholl. Principal funding for this work came from the European Research Council (ERC Advanced grant (SEE-MAGIC) grant no. 101098056) to J.O.K., with additional support coming from an ERC Consolidator grant (MOSAIC; no. 773026) to J.O.K, the Volkswagen Foundation (VW-95826) to J.O.K, and from EMBL core funding. M.R.C. received support from the EMBL Interdisciplinary Postdoc (EIPOD4) program 4 under Marie Sklodowska Curie Actions COFUND (grant agreement no. 847543), enabling interdisciplinary studies in the Pepperkok and Korbel groups. We acknowledge the EMBL core facilities and services for support in high-performance computing (IT), sequencing (GeneCore), chemical synthesis (Chemical Biology), imaging (Advanced light microscopy) and cell sorting (FACS).

## Data availability

All genomics data generated in this study (Strand-seq, as well as short and long-read bulk WGS) are available at ENA under the following accession: PRJEB78885. We re-analysed publicly available data from the PCAWG^8^ and TCGA resources to compare our findings to those previously made in cancer genomes. The raw WGS data generated by TCGA can be accessed through controlled data access application via dbGAP under study accession code phs000178.

## Code availability

The software automation components and step-by-step instructions for running MAGIC experiments are available in the magic_automation repository (https://git.embl.de/cosenza/magic_automation). For image analysis, computer vision, and image processing, visit the magic_tools repository (https://git.embl.de/cosenza/magic_tools). Additionally, tools for analysing Strand-seq data and single-cell copy number calling are provided in the strandtools repository (https://git.embl.de/cosenza/strandtools). The script for estimating basal CA rates is provided at https://git.embl.de/cosenza/ca_rates_estimation. The code used in this study to analyse WGS data is available at: https://github.com/cortes-ciriano-lab/osteosarcoma_evolution.

## Author contributions

M.R.C., J.O.K. and R.P. conceived the project, with J.O.K. providing scientific direction and supervision. M.R.C. and A.H. developed the microscope automation software and integration with AutoMicTools. M.R.C. developed *magic_tools*. M.R.C. designed and performed the photolabeling optimization experiments. M.R.C. and A.G. designed and performed the long-term live-cell imaging experiments, M.R.C., A.G. and N.L.S. analysed the image data. N.L.S. performed Western blotting experiments. M.R.C., A.G., N.L.S. designed and performed MAGIC experiments with the support from P.H.. A.G. performed single-cell clone propagation experiments, low-pass WGS and long-read sequencing characterization of clones. Long-read data was analysed by A.G. and T.R.. M.R.C. designed the FISH experiments, which were performed by A.G.. M.A.J. developed the convolutional neural network. P.H. prepared Strand-seq libraries. Single-cell data was analysed by M.R.C., A.G., N.L.S., A.A.D. and J.O.K.. A.A.D. designed and performed copy-number pattern analysis in PCAWG data, with support from M.R.C.. A.A.D. and S.Z.D. designed and performed the isochromosome analysis in primary cancer genomes, with guidance from J.O.K. and I.C.C.. B.E.U. designed and tested the CRISPR guides, under guidance of A.K.. CRISPR experiments were performed by B.E.U. and A.G.. M.R.C. developed the agent-based statistical model with support from E.G.. M.R.C. and J.O.K. wrote the core of the manuscript, with contributions from all authors. J.O.K., R.P., A.K., T.R., A.G., N.S., A.A and I.C.C. critically reviewed and edited the manuscript.

## Ethics declarations

The following author has previously disclosed a patent application (no. EP19169090) that is relevant to the use of Strand-seq for somatic structural variation analysis: J.O.K. The remaining authors declare no competing interests.

## Methods

### Statistical analysis

Unless otherwise stated, we employed the following system to indicate significance levels in the figure panels: * *P* < 0.05; ** *P* < 0.01; *** *P* < 0.001. Statistical tests used are indicated in the main text or figure caption, with specific tests for chromosome biases and breakpoint as well as SCE locations detailed in the **Supplementary Methods**.

### Cell culture and cell line development

MCF10A (CRL-10317, American Type Culture Collection) and RPE-1 (CRL-4000, American Type Culture Collection) cell line and their *TP53-/-* derivatives were cultured at 37°C with 5% CO2 atmosphere and 100% humidity, in DMEM/F12 medium (1:1) without phenol red (Gibco), supplemented as follows: RPE-1 medium was further supplemented with 10% FCS and antibiotics; MCF10A medium with 5% horse serum (Thermo Fisher Scientific), 20 ng/ml human EGF (Biotrend), 0.5 mg/ml hydrocortisone (Sigma-Aldrich), 100 ng/ml cholera toxin (Sigma-Aldrich), 10 μg/ml recombinant human insulin (Sigma-Aldrich) and antibiotics. MCF10A *TP53-/-* cells have been kindly provided by Prof. Dr. Claudia Scholl (Laboratory of Applied Functional Genomics, DKFZ), while RPE-1 *TP53-/-* variants were generated in a previous study from our lab^56^. All cell lines tested negative for mycoplasma contamination.

A plasmid carrying H2B-Dendra2^82^ (Addgene, plasmid #75283) was introduced in all cell lines by transfection: 20,000 cells were seeded in a glass-bottom slide (Nunc LabTek 8-well) and transfected with 20µL of transfection mixture at 4:1 ratio of Fugene HD (Promega) to DNA in Optimem (Thermo Fisher Scientific). Transfection success was assessed 48 hours later by fluorescence microscopy, cells were transferred into two 10 cm dishes and G418 antibiotic was added at 200 µg/mL (MCF10A) or 400 µg/mL (RPE-1) for selection. Two weeks later, well separated, fluorescent colonies were visible and were isolated by pipetting, transferred to 24-well plates and grown into stable cell lines. Stable-transfectants for RPE-1 WT and RPE-1 *TP53-/-* were instead collected in pool and isolated by single-cell sorting using a BD FACSAria at 1.0 flow rate, with a 130um nozzle, dispensed in a flat-bottom, 96-well plate (ThermoFisher, Nunc plates) with normal growth medium.

### MAGIC: Autonomous Platform for *de novo* CA formation studies

MAGIC leverages machine learning and automated microscopy to perform targeted photolabeling of cells-of-interest, for subsequent fluorescence-activated cell sorting and downstream analysis. A MAGIC experiment comprises three phases: (1) The preparation phase, where cells are seeded and additional treatments can take place, such as targeted DSB induction or staining by DACT-1. (2) The photolabeling phase, where targeted illumination^79,80,83^ takes place using automated microscopy. (3) The harvest phase, when cells are collected and isolated by FACS. These steps are outlined below, accompanied by further details presented in the **Supplementary Methods**.

#### Preparation

During this phase cells are prepared to undergo the targeted photolabeling procedure. Additional treatments can take place, such as targeted DSB induction, staining with live-cell dyes and adding BrdU for Strand-seq. To enable photolabeling, we engineered MCF10A and RPE-1 cell line models to constitutively express H2B-Dendra2, a monomeric fluorescent protein that undergoes irreversible photoconversion with 405 nm light, which also enables the visualisation of nuclear atypia without impacting mitotic fidelity^84^. As an alternative, for RPE-1 wild-type cells we also utilised DACT-1, a photo-activatable cell tracking dye, that converts to a bright red-fluorescent state upon 405 nm light exposure, further details are available in the **Supplementary Methods**.

Cells were seeded in up to 4 wells of a µ-slide 8-well dishes (Ibidi). Seeding density was adjusted to have about 40,000 cells one day before experiment start. In the case of Strand-seq downstream analysis, BrdU (40 µM) was added to the cells prior to the start of photolabeling. One control slide, without BrdU, was also prepared to adjust gating strategies during single-cell sorting. In the case of targeted DSB-induction experiments, RNP-complexes were delivered by electroporation 48 hours before the start of the experiment and up to two different sgRNA were examined during a single experiment. Further details can be found further below.

#### Photolabeling

Living cells are then transferred on an LSM 900 microscope (Zeiss) with confocal and widefield imaging capabilities, and an environmental chamber with temperature and CO2 control. MAGIC relies on full microscope automation and computer vision for laser-assisted, phenotype-driven targeted illumination of single-cells at scale. The system includes three software components: a microscope control script, an image analysis manager based on *AutoMicTools*, and a python package, *magic_tools*, which we designed for advanced image processing.

The microscope control script automates autofocusing, micronuclei identification, and photoconversion of target nuclei across several positions. Autofocus is achieved by detecting the glass bottom dish reflection using a 639nm laser and *AutoMicTools* analysis. For micronuclei identification, a Z-stack image centred on the focused slice is analysed on an image analysis server driven by *magic_tools*. Photoconversion involves using micronuclei coordinates to define ROIs of the corresponding parental nuclei, which are then selectively illuminated with a 405 nm laser. Pre- and post-experiment images are acquired before the microscope moves to the next position.

We run the photolabeling experiment overnight and up to 24 hours, to achieve a yield of 700 to 2000 phololabeled cells, depending on the experimental conditions. A detailed description of the automation software and the image analysis pipeline can be found in **Supplementary Methods**.

#### Harvesting

Following photolabeling, cells were harvested and target cells were isolated by single-cell sorting. In case of Strand-seq experiments, at the end of the photolabeling phase, cells were stained for one hour with Hoechst 33342. Cells were harvested with 0.25% Trypsin (Gibco) and resuspended in buffer (8% FBS in 1X PBS, supplemented with Hoechst 33342 5μg/ml and BrdU 40 µM). Single cells were sorted using a BD FACSAria, in purity mode, with a 100 or 130µm nozzle and dispensed lysis buffer or fresh medium in a flat-bottom, 96-well plate (ThermoFisher, Nunc plates). We employed the following gating strategy: we selected first the general population in forward and side scatter and we excluded doublets. Then, cells were sub-gated for photolabeled cells as shown in **Figure 1G** for H2B-Dendra2 or **Figure S2G** for DACT-1. When employing Strand-seq, the singlet population was further filtered to select cells with a quenced Hoechst signal that had thus incorporated BrdU^85^. Cells harvested from control slides were used to optimally adjust gates to exclude false positives.

### Long-term live-cell imaging

The live-imaging experiment for nuclear and mitotic phenotype^24,86^ scoring was carried out over the course of 72 hours. MCF10A cells stably expressing H2B-Dendra2 were seeded at a 15-25% confluence on µ-slide 8-well dishes (catalogue no.: 80806; Ibidi), and images were acquired every 10 minutes with a Plan-Apochromat 20X/0.8 M27 air objective using the LSM900 confocal microscope (Zeiss). Manual annotation was performed with the assistance of a customised tool written in python. Mitotic phenotype and nuclear morphology for parental cells and the first generation of daughter cells were annotated, as described in **Figure 1B**.

### Optimization of photolabeling parameters

MCF10A cells stably expressing H2B-Dendra2 were seeded on μ-slides (Ibidi) and imaged on an LSM900 confocal microscope (Zeiss). To determine Dendra2 photoconversion dynamics, we performed 5 bleaching rounds, each with 10 laser-scanning iterations with a 20X objective and 405 nm laser, at scanning speed 8 and power at 0.5% in the low-intensity power range. Images in green and red channels were acquired at the beginning and end of each round. The fluorescence intensity of ten photoconverted nuclei and five non-photoconverted control nuclei per field of view was quantified on manually defined ROIs with ImageJ. Data was then processed and analysed with custom python scripts.

To assess phototoxicity from targeted illumination, MCF10A and RPE-1 cells seeded on μ-slides (Ibidi) were photoconverted with settings used in the MAGIC pipeline and followed by confocal microscopy. Images for native and photoconverted Dendra2 fluorescence channels were acquired with a 20X objective over the course of 24 hours. Cells were tracked manually and their fate annotated. No cell death was detected for the photoconverted cells within the analysed timeframe.

### Single-cell genomic sequencing with Strand-seq

For Strand-seq, an intermediate coverage, whole genome amplification free single-cell method^38–40,87^, we performed cell sorting as in the original procedure^38^ with important adjustments to accept whole cells as input, in order to avoid loss of cytoplasmic DNA material and micronuclei during nuclei isolation. Cells were incubated with Hoechst 33342 (5μg/ml) for 60 minutes, as it is cell-membrane permeable. Then cells were harvested with 0.25% trypsin (Gibco) and resuspended in buffer (8% FBS in 1X PBS, supplemented with Hoechst 33342 5μg/ml and BrdU 40 µM). Single cells were sorted using a BD FACSAria, in purity mode, with a 100 or 130µm nozzle and dispensed in a flat-bottom, 96-well plate (ThermoFisher, Nunc plates), with freeze buffer supplemented with 0.2% NP-40 (Thermo Fisher Scientific) to ensure membrane lysis and DNA accessibility in subsequent protocol steps. Strand-seq libraries were prepared at large-scale using a liquid handling robotic platform as described previously^39^. Libraries were sequenced on a NextSeq5000 (MID-mode, 75 bp paired-end) followed by demultiplexing. Reads were aligned to GRCh38 reference assembly with BWA-MEM version 0.7.17, yielding a median of ∼285,000 mapped unique fragments per cell, and further processed with MosaiCatcher^39,88^ and *strandtools* (see below).

### Single-cell *de novo* CA discovery and classification

We discovered a wide variety of *de novo* CA classes leading to chromosomal or segmental copy-number imbalances by integrating read coverage and Watson(W)/Crick(C) template ratios^39^, enabling high-resolution CA calling in Strand-seq data. Extending the functionality of the previously released MosaiCatcher tool^39^, we designed *strandtools*, which is tailored for the specific task of handling *de novo* CA discovery in single cells under diverse ploidy backgrounds (further details, see **Supplementary Methods**). To achieve high confidence CA classification, we integrated read depth, strand orientation and haplotype information in each cell^39^, to characterise segmental alterations and assign them to one of the following CA classes: chromosome loss, chromosome gain, interstitial loss, interstitial gain, terminal loss, terminal gain, terminal multi-step, complex CA and chromothripsis (a complex CA subclass). Chromosome gains and losses affect a whole chromosome, from p-ter telomere to q-ter telomere. Interstitial gains and losses are isolated CAs between two breakpoints, within one chromosome arm. As terminal alterations, we refer to all CAs that involve a portion of a chromosome, from a breakpoint anywhere along a chromosome arm to the telomere of that same arm. Therefore, terminal gains and losses are simple CAs, with one isolated, altered segment spanning from a breakpoint to the telomere of one chromosome arm. Terminal gains are annotated as inverted duplications if the gained segment is in opposite strand orientation compared to that of the original homolog with the same haplotype^39^. Terminal multi-step CA are a sequential combination of gains and losses that are affecting the terminal portion of a chromosome arm. The terminal multi-step class also includes all cases of localised oscillations arising alongside terminal gains and losses.

Complex CAs are defined as events that include more than 2 breakpoints, can affect either one or both arms of the same homolog and can be composed of non-adjacent, altered segments. As such, complex CAs cannot be resolved as terminal multi-step. Chromothripsis events extending over large chromosomal regions, such as a chromosome arm, are included under the complex CA class. These events show characteristic copy-number oscillation between typically two copy-number states, affecting one single haplotype and with oscillating segments allowed in either strand orientation^39,41^. With regards to experiments on targeted DSB induction along chromosome arms, we likewise considered all copy-number imbalanced CA classes. In addition, we specified whole arm alterations in the case of isolated gains and losses affecting more than 90% of a chromosome arm, and amplifications in case of isolated gains with a copy-number increment of 2 or more compared to the baseline. All single-cell CA annotations are available in **Supplementary Tables 7, 8 and 9**.

### Targeted induction of DSBs

CRISPR components, designed as described in the **Supplementary Methods**, were delivered in the form of ribonucleoprotein (RNP) complex via Neon Electric Transfection System (10 µl kit; catalogue no.: MPK1096; Thermo Fisher). First, the RNP complex was formed by incubating 0.3 µl of Alt-R® S.p. Cas9 Nuclease (catalogue no.: 1081059; IDT) with 0.2 µl Resuspension Buffer R (Neon 10 µl kit) and 1 µl of designed sgRNA for 20 min. Cells (0.5e6 per reaction) were prepared for electroporation as described in the manufacturer’s manual. Concentration of Cas9 nuclease in the final RNP/cell suspension was 1.5 µM, and that of sgRNA was 3.6 µM. Electroporation parameters of 1400 V, 20 ms, 2 pulses were used for both RPE-1 and MCF10A cells. Transfected cells were diluted in antibiotics-free cell culture medium and different amounts (between 36.000-72.000) were seeded into four central wells of µ-slides containing 300 µl of antibiotics-free medium. The medium was replaced by BrdU (40 µM) containing fresh medium 48 h post-transfection, and the slide was immediately transferred into the confocal microscope for imaging.

### Clone generation from single-cells

Cells were subjected to automated photolabeling, harvested with 0.25% trypsin (Gibco) and resuspended in buffer (8% FBS in 1X PBS). Single cells were sorted using a BD FACSAria at 1.0 flow rate, with a 130um nozzle to minimise cell damage, and dispensed in a flat-bottom, 96-well plate (ThermoFisher, Nunc plates) with normal growth medium. Formation of viable colonies was visually assessed daily with a phase-contrast microscope from day 7 to 14 post sorting. At the 2 week mark, clones were transferred to 6-well plates, and grown to confluence to be frozen for future experiments and prepared for sequencing.

### Low-pass short-read WGS of clones

27 MCF10A cell pellets (18 clones deriving from micronucleated cells, 9 control clones) were subjected to low-pass Illumina sequencing (NextSeq2000, P3, 100bp paired-end sequencing) at EMBL’s Genomics Core Facility, to an approximate genomic coverage of 1× for screening purposes. Reads were aligned to the GRCh38 genome reference with BWA^89^, and read depth based CA calling was performed with support of the Control-FREEC tool^90^.

### Long-read WGS of clone 7

Clone 7 was re-established to obtain 10*10^6 cells for ONT sequencing. The library was prepared using the SQK-LSK114 ligation kit, and sequencing performed on PromethION flow-cells at the German Cancer Research Center. The obtained coverage was 16×, and the reads showed an estimated N50 of 13.97kb. Reads were aligned to the GRCh38 genome reference with minimap2.^91^ SV calling was performed with Sniffles^92^ and Delly^93^, and calls were manually curated to exclude false positives. Haplotype phasing of the ONT reads was performed with Whatshap^94^.

### Modelling *de novo* CA rates

We developed an agent-based model^95^ to simulate CA acquisition in a growing population of cells, considering mitotic errors and micronuclei generation. During the simulation, cell agents are allowed to move between the states depicted in **Fig. 3F**. The probability *p_ij_* of transitioning from state *i* to state *j* is derived from long-term live-cell imaging experiments. Each cell agent is designed to possess three main attributes: cell-cycle status, micronucleus status and CA status. The micronucleus status captures whether the cell possesses a micronucleus or not. The cell-cycle status keeps track of an internal clock that simulates advancing through cell cycle until mitosis. Cell cycle duration is set at the median cell cycle duration measured in imaging experiments. The CA status captures whether the cell possesses a *de novo* CA. When the internal cell-cycle clock reaches the end, mitosis or arrest occurs: the cell agent can move from interphase to a mitosis state (normal, laggard or bridge) or arrest. To simulate cell division, the current agent is moved to the arrest state and two new cells are generated and assigned to state 1 or 5, according to the transition probability associated with that specific mitosis type. Moreover, during mitosis, each cell has the possibility of acquiring a *de novo* CA according to the assigned rate *R*. Arrested cells are then removed from the simulation. Each simulation is initiated with an initial population of 50 cells and it is stopped when the population reaches size 50,000, as we empirically found that the micronuclei and CA frequency usually stabilise by this time. At the end of the simulation we compute the sum of squared error between the simulated and target *de novo* CA frequencies. Details on how bound-constrained minimisation was employed to the CA rate estimation are the **Supplementary Methods**.

### Fluorescent *in-situ* hybridization

MCF10A cells were seeded on coverglass slides, subjected to targeted DSB induction, and allowed to recover for 48 hours. Metaphase spreads were then prepared *in situ* directly on coverslips, as described elsewhere^96^. Sub-centromeric probes for chromosome 7 p and q arms were purchased from KromaTiD (Biocat catalogue no.: CEP-0013-C-KTD, CEP-0014-A-KTD). FISH was performed according to manufacturer instructions. After post-hybridization washes, DNA was stained with Hoechst 33342 and slides were mounted in anti-fade medium (Vectashield, Vector Laboratories). FISH images were acquired on a LSM900 confocal microscope (Zeiss) at 40X magnification and signals evaluated visually.

