## Supplementary Information for "Origins of *de novo* chromosome rearrangements unveiled by coupled imaging and genomics"


**SUPPLEMENTARY INFORMATION** **1**
**Supplementary Figures** **2**
**Supplementary Tables** **20**
**Supplementary Methods** **21**
Photolabeling strategies 21
Microscope automation and imaging 21
Online image analysis with magic_tools 22
Strandtools – an optimised single-cell CA calling algorithm for Strand-seq data 23
Computational sister cell pair discovery from Strand-seq data 24
XGBoost classifiers training 24
Convolutional neural network based micronucleus classifier training 24
Designing single-guide RNA 24
Statistical testing for biases in CA frequencies, SCEs and breakpoint locations 25
CA rate estimation by bound-constrained minimization 26
Western blotting 26
**Supplementary Notes** **27**
Pan-cancer WGS cohorts 27
Comparing copy-number features from spontaneous micronuclei with the PCAWG resource 27
Quantification of aneuploidies in the PCAWG resource 29
Inference of isochromosomes using bulk WGS data 29
**References** **31**

### Supplementary Figures

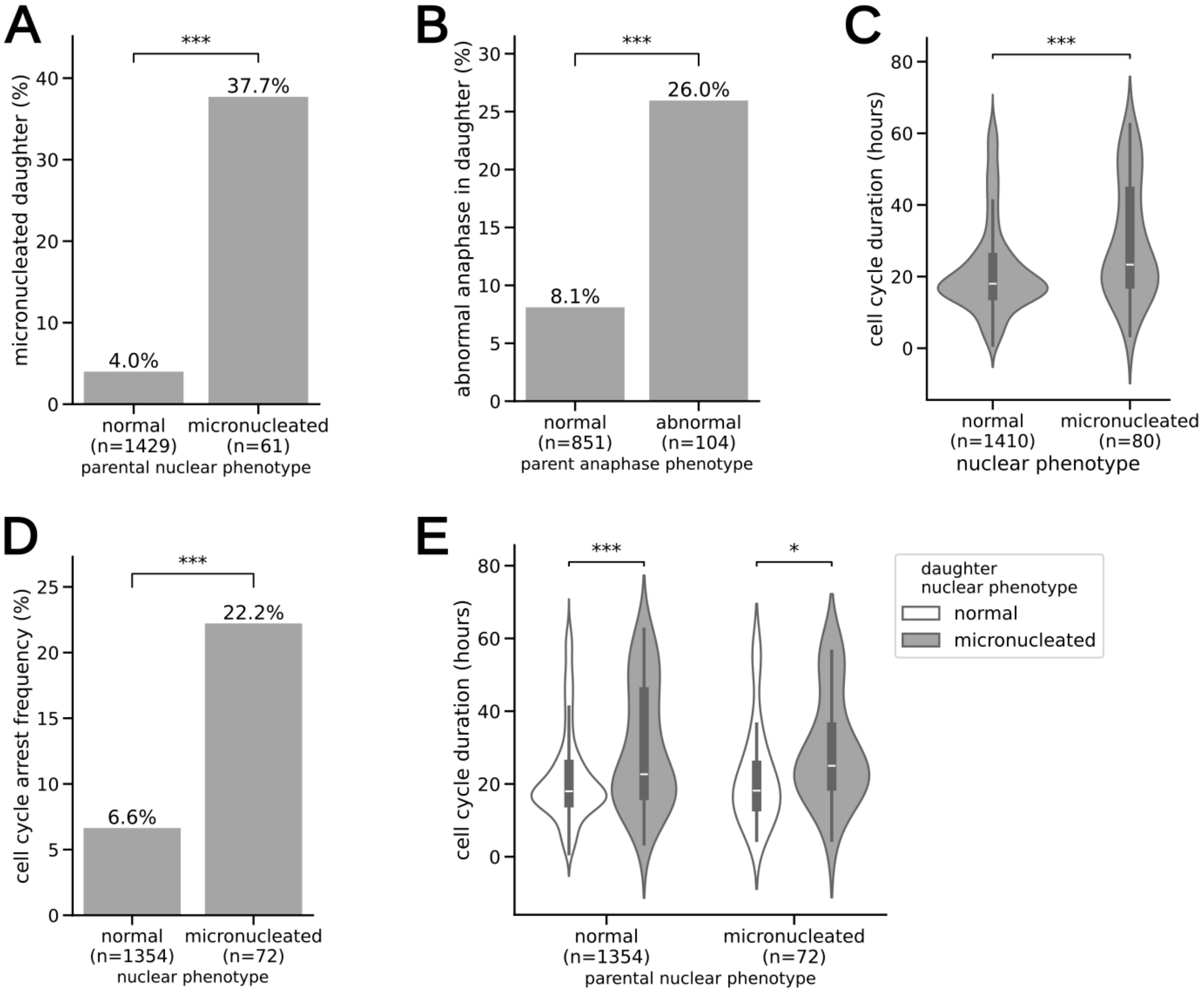


**Figure S1 | Long-term live-cell imaging analysis in MCF10A cells**
*(A) frequency of micronucleated parent cells that generate at least one micronucleated daughter (Fisher's exact test, P < 0.001) (B) frequency of abnormal anaphase in daughter cells, depending on the preceding anaphase phenotype of the in parent cell (Fisher's exact test, P < 0.001) (C) cell cycle duration in hours for daughter cell generation. Duration measured from mitosis to mitosis (Mann-Whitney U test, two-sided, P < 0.001) (D) Frequency of cell cycle arrest, defined as cells with a cell cycle duration longer than the 99th percentile (Fisher's exact test P < 0.001) (E) Cell cycle duration of daughter cells depending on the nuclear phenotype seen in the parental generation (Mann-Whitney U test, two-sided, normal P < 0.001, micronucleated P < 0.05). (for significance levels see* ***Methods****, * P<0.05, **P<0.01, ***P<0.001)*

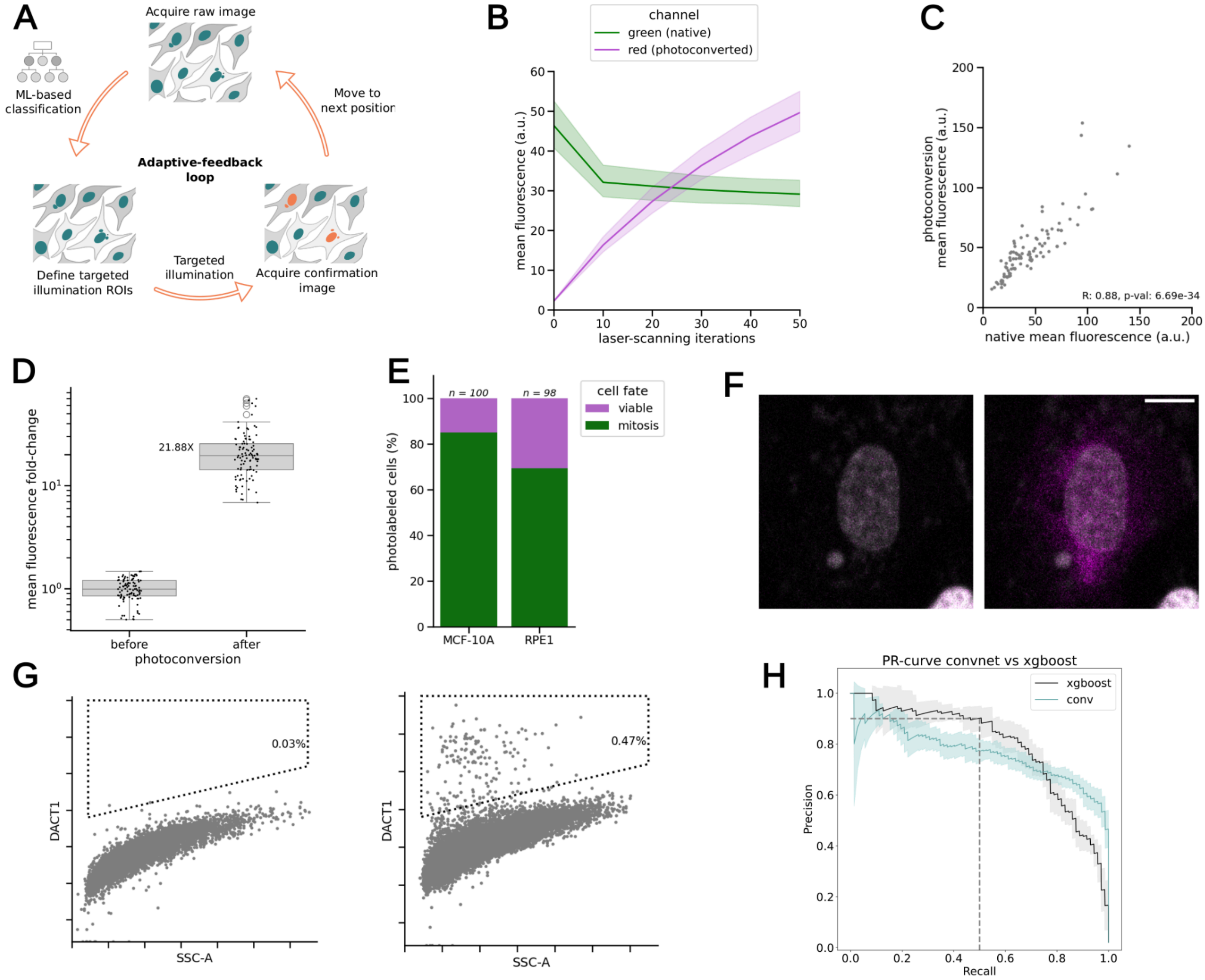


**Figure S2 | Establishing imaging parameters for setting up MAGIC**

*(A) Schematics depicting the adaptive-feedback loop at the core of MAGIC's microscope automation. (B) Photoconversion dynamics of targeted illumination of H2B-Dendra2. Mean fluorescence was measured in 151 cells every 10 laser-scanning iterations. Color band, 95% confidence interval. (C) Correlation between native (green) and photoconversion (red) mean fluorescence of photolabeled nuclei in MCF10A cells expressing H2B-Dendra2 (Pearson correlation coefficient) (D) Fold change in mean nucleus fluorescence before and after photolabeling in MCF10A cells expressing H2B-Dendra2. Center line, median; box limits, upper and lower quartiles; whiskers, 1.5x interquartile range; points, outliers. (E) Fate of imaged cells followed for 24 hours after photolabeling (F) RPE-1 cells before (left) and (after) photoactivation DACT-1 dye (magenta). Nuclear DNA (grey) visualised by NucSpot 488. Scale bar: 10 µm (G) FACS profile of MCF10A H2B-Dendra2 cells in control (left) and photolabeled (right) conditions. Side scatter signal area (SSC-A) versus DACT-1 fluorescence is plotted, and the sorting gate is represented by a dashed line. (H) Precision-recall curves for XGBoost (grey) and convolutional (conv, cyan) neural network models for micronuclei classification. Coloured bar represents standard deviation of performance on five different training-test dataset splits.*

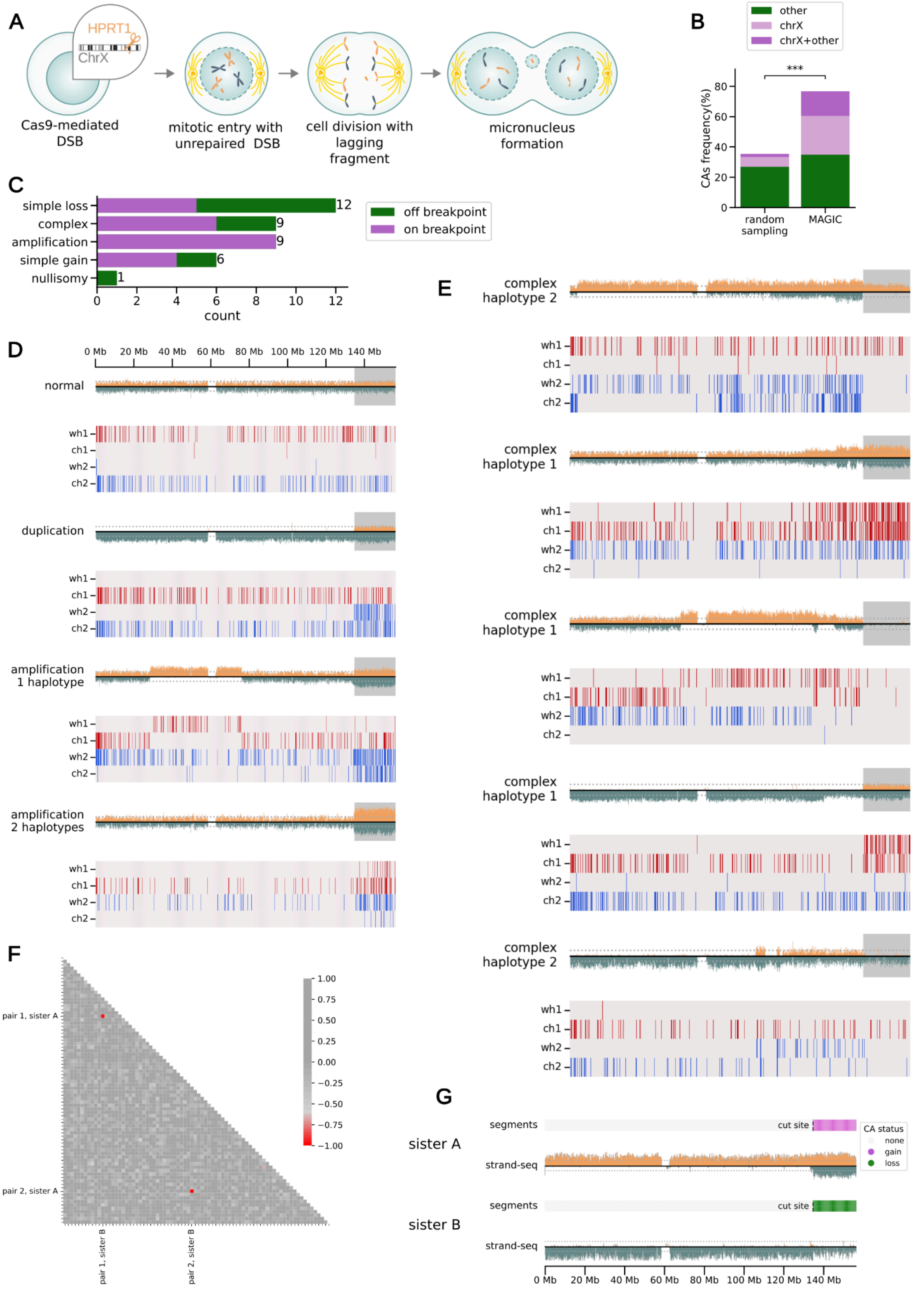


**Figure S3 | Verification of the MAGIC platform through Cas9-induced CAs**

*(A) Schematic showing how Cas-9 mediated DSB can lead to micronucleus formation. In this example Cas-9 cuts at the HPRT1 locus on chromosome X in G2 phase, generating an acentric chromosome*

*fragment lacking the centromere. If the cell enters mitosis without repairing the DSB, the acentric fragment cannot participate in mitosis as it would be missing the kinetochore necessary for spindle orientation. As a consequence, the acentric fragment would lag behind at anaphase and be inherited by either daughter cell, forming a micronucleus. (B) Frequency of CAs in single cells randomly sampled, or selected using MAGIC. chrX: chromosome X (Fisher's exact test, P<0.001) (C) Breakdown of copy number alterations after targeted DSB induction at the HPRT1 locus. Simple loss and simple gain indicate an isolated change in copy number. Amplification refers to all isolated copy number gains of two or more. Nullisomy is the isolated and complete loss of the chromosome segment. Complex CAs are defined as events that involve more than 2 breakpoints, which can comprise combinations of copy number gains and/or losses. (D,E) Strand-seq based haplotype-aware analysis[1] of acentric fragment copy number gains and amplifications of the cut-off segment (D) and complex CA affecting a single haplotype (E). Grey background in strand-seq plots: cut-off fragment. Phased haplotags[1] are represented by red or blue lines for haplotype 1 and 2, respectively. wh1: watson strand, haplotype 1. wh2: watson strand, haplotype 2. ch1: crick strand, haplotype 1. ch2: crick strand, haplotype 2 (F) Strand orientation state correlation between single-cells as a predictor for sister cell pair relationship. Two sister cell pairs displaying a pronounced negative correlation are shown in red (Pearson correlation coefficient; scale bar shown to the right). (G) Example of a reciprocal CA in sister cells. Sister A carries a copy number gain of the cut-off fragment generated by the cut (top), while sister B  carries a copy number loss for the same segment (bottom). Cut site annotated by dashed line.*

---

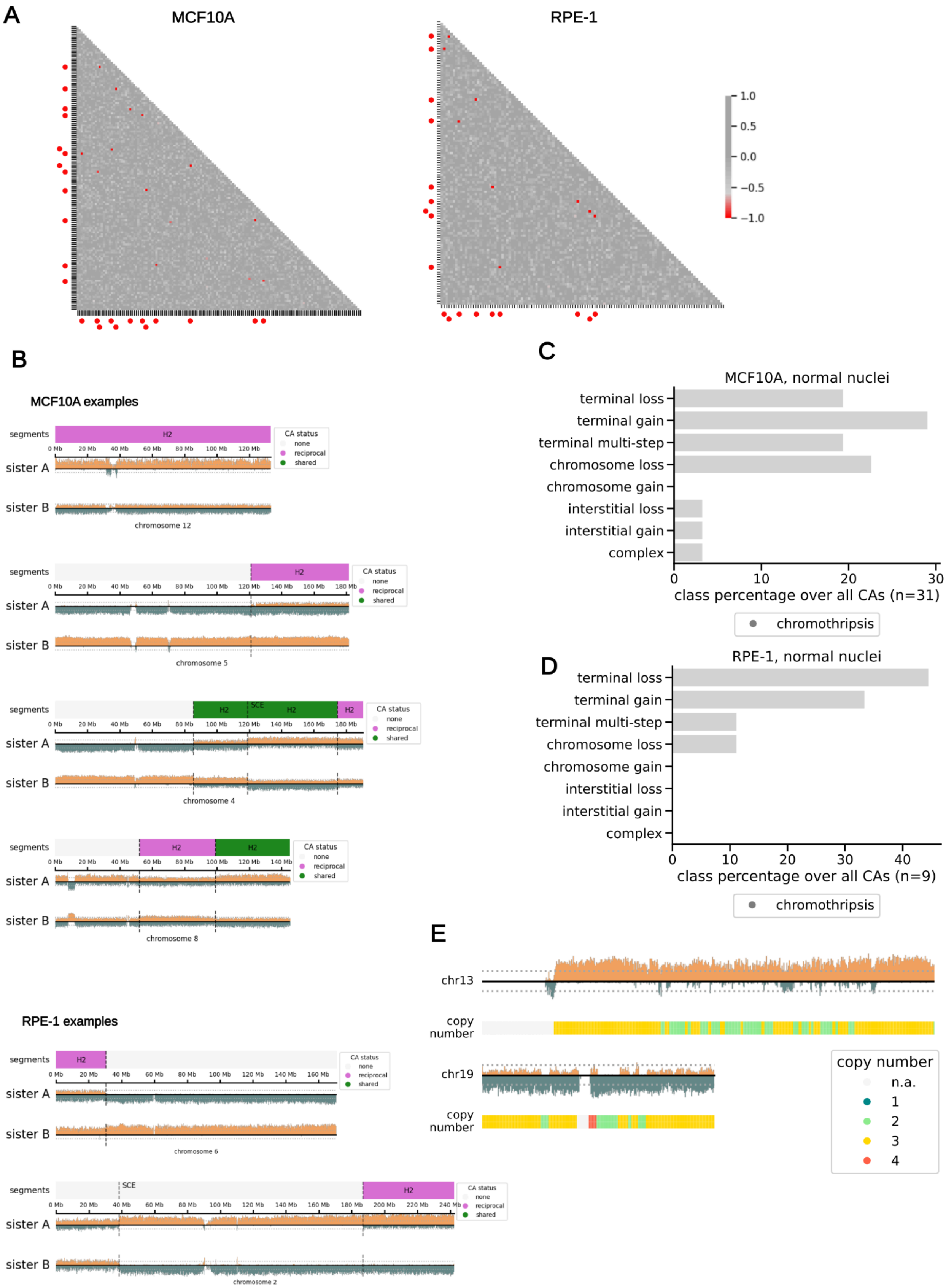


**Figure S4 | Sister cell analysis and CA examples in near-diploid non-transformed cells**

*(A) Sister cell strand-state anti-correlation analysis for MCF10A (left) and RPE-1 (right) micronucleated cells (Pearson correlation coefficient) – revealing several sister cell pairs through single-cell genomic analysis. Sister cell pairs are indicated by red circles on the heatmap axis. (B)*

*Reciprocal CA examples, detected using Strand-seq[1], in MCF10A and RPE-1 cells, with Annotated segment boundaries marked by dashed lines. (C,D) Breakdown of CA classes in normal MCF10A (D) and RPE-1 (E) cells, with the class percentage shown relative to all CAs. (E) Chromothripsis examples from spontaneous micronuclei in MCF10A cells*

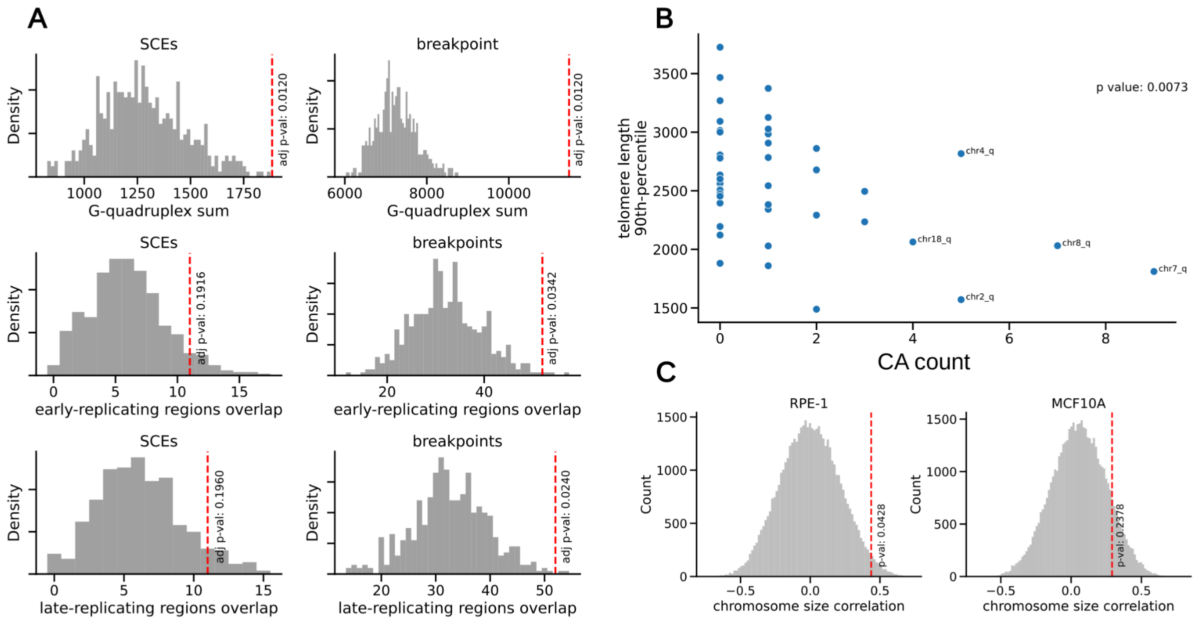


**Figure S5 | Breakpoint features of CAs from spontaneous micronuclei**

*(A) Distribution derived from permutation of breakpoint location against G-quadruplex sites (top row), early (middle row) and late (bottom row) replicating regions. Red dashed line: observed statistic (see* ***Methods****). The analysis was conducted using MCF10A cells, from sporadically micronucleated cells. (B) Correlation between CA count and ONT-sequencing based telomere length per chromosome arm, expressed as 90th percentile of read length (Pearson correlation; see* ***Methods****). (C) Distribution derived from permutation of determined CA numbers in spontaneously micronucleated cells against chromosome size. Red dashed line: observed statistic.*

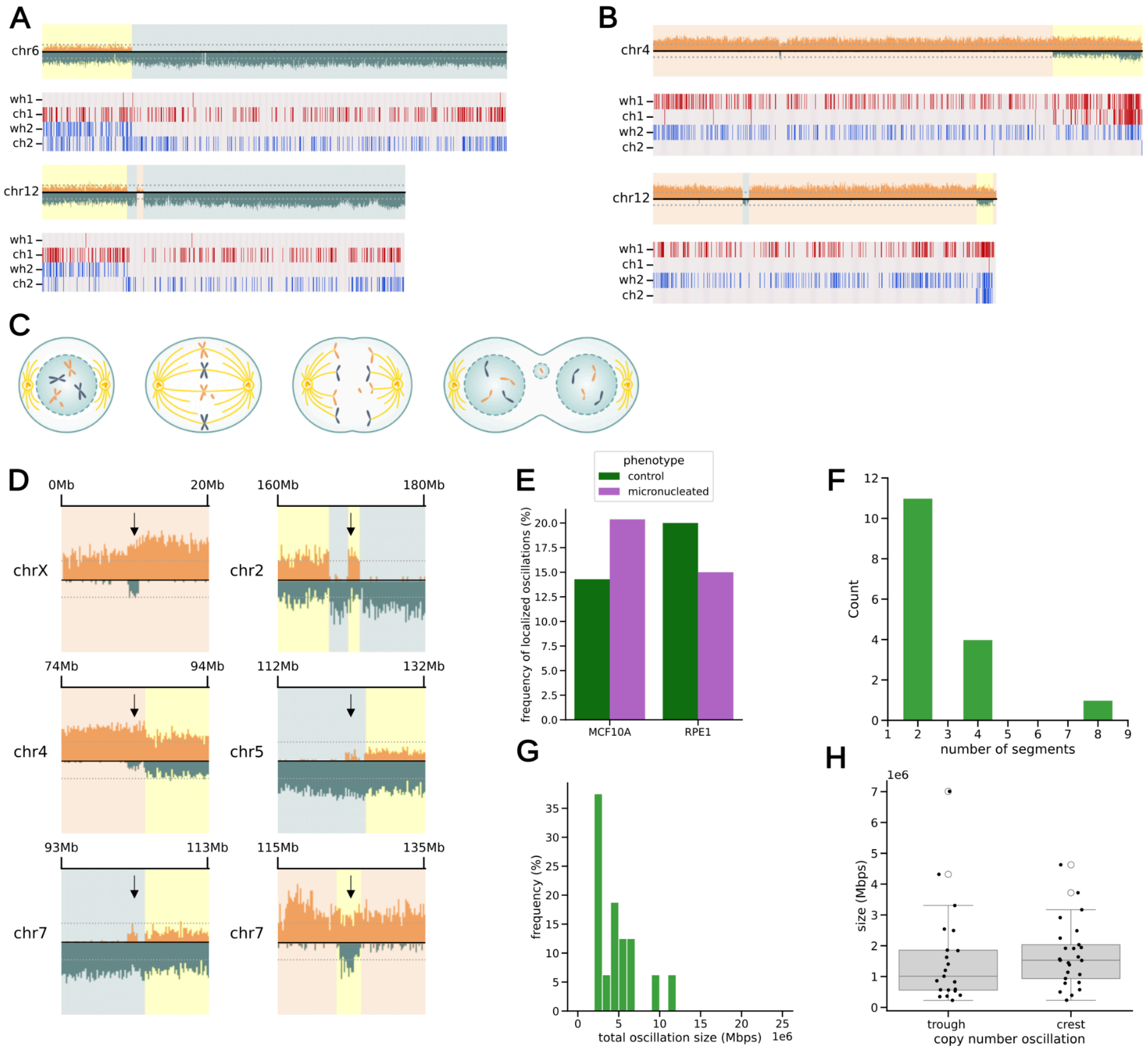


**Figure S6 | Analysis of sporadic bridge-mediated CAs**

*(A,B) Strand-seq based haplotype-resolved examples of terminal inverted duplications (B) and terminal multi-step alterations (B). For each example, we depict at the top: strand-seq plot, bottom: haplotag[1] localisation. In (A) inverted terminal duplications is characterised by the presence of haplotype 2 (H2) haplotags on the W strand for both chromosome 6 and 12 examples. In the (B) upper panel, chromosome 4 carries two adjacent copy number gains, affecting haplotype 1 (H1) and both strands. In the lower panel, chromosome 12 carries one inverted duplication on H2, with an adjacent terminal deletion of the sample haplotype. (C) Scheme showing mitosis entry with unrepaired DSB leading to micronucleus formation. (D) Localised copy-number oscillation examples indicative for small to intermediate scale complex SVs. Arrow: oscillation peak. (E) Frequency of the localised oscillation pattern in MCF10A and RPE-1 cells. (F,G,H) Features of the localised oscillation pattern: number of segments composing the oscillation (F), overall oscillation (G), trough and crest (H) oscillation size. The data depicted includes cases from both WT and TP53-/- cell line models. Center line, median; box limits, upper and lower quartiles; whiskers, 1.5x interquartile range; points, outliers.*

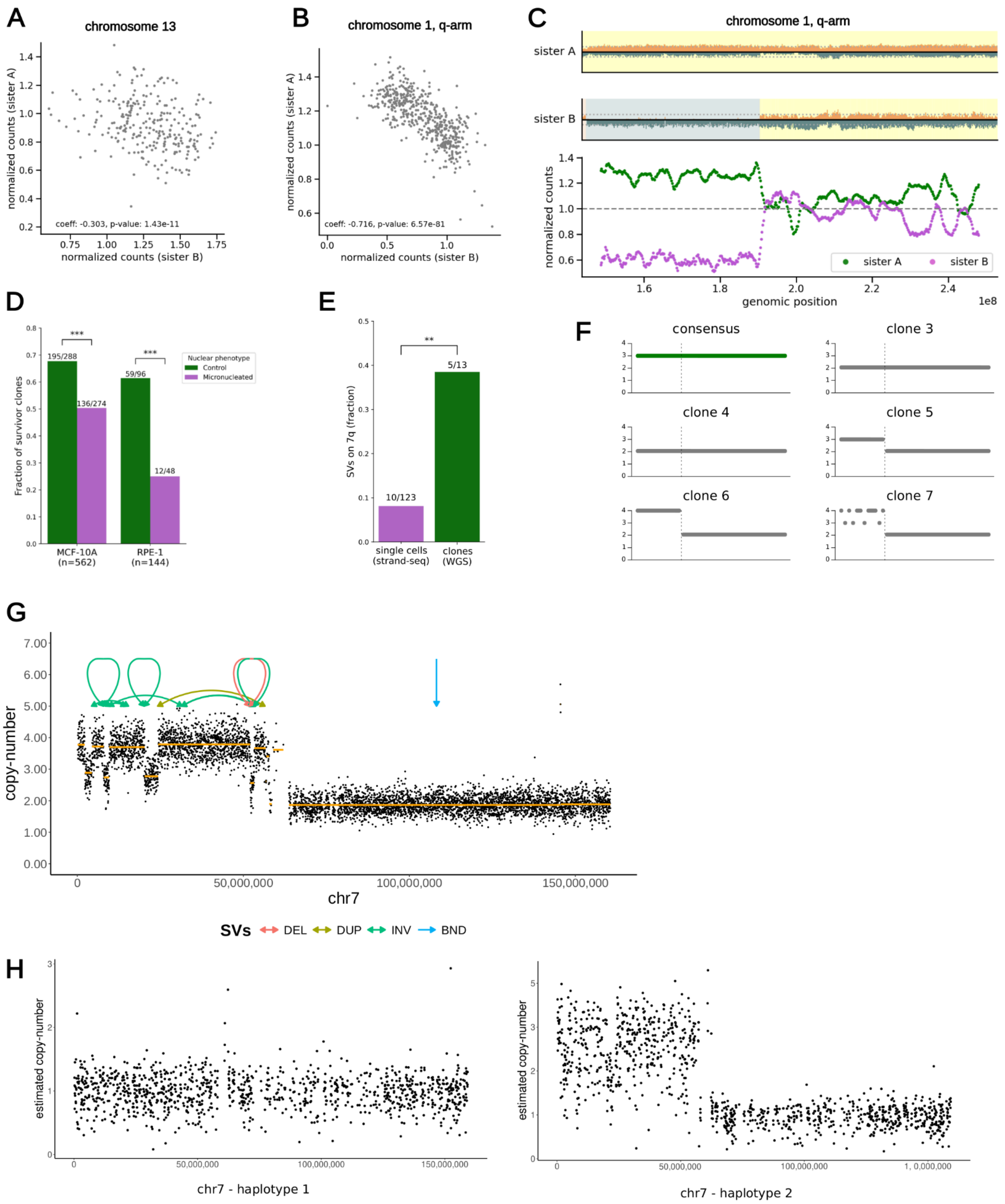


**Figure S7 | Analysis of chromosome-scale chromothripsis, and clonal propagation experiments**
*(A,B) Sister cell read-count anti-correlation for chromosome 13 (A) and chromosome 1 q-arm (B) (Pearson correlation coefficient), verifying the reciprocal segregation of shattered DNA fragments. (C) Chromothripsis in a sister cell pair, affecting chromosome 1, q-arm, determined in a complex ploidy background. Top, Strand-seq plots with oscillatory copy number pattern. Bottom, smoothened and normalised counts along chromosomal positions. (D) Fraction of surviving clones after MAGIC isolation. (E) 7q loss enrichment in propagated clones. (F) Schematic of 7q hits in micronucleated clones, which include 7q-arm loss, chromosome 7 losses, and isochromosomes resulting in 7q-loss. (G) ONT long-read sequencing based copy number plot of chromosome 7 for the micronucleated clone 7; arrows on top represent the boundaries of the different classes of SVs identified by both Delly[2] and Sniffles[3] (DEL: deletion; DUP: duplication; INS: insertion; INV: inversion; BND: translocation). The chromosome presents a duplication of the p-arm coupled to the deletion of the q-arm, indicating isochromosome formation; in addition, a chromothripsis event is identified based on*

*the presence of an oscillating copy-number pattern (in this case, between copy-number 3 (CN3) and CN4) and on the simultaneous occurrence of multiple SVs on the affected chromosome arm consistent with randomness of DNA fragment joins[4]. As a consequence of isochromosome formation followed by chromothripsis, three copy-number states are seen across the chromosome 7 genomic coordinates. (H) Copy number plot of chromosome 7 of the micronucleated clone 7, resolved by haplotype. The haplotype phasing was performed with Whatshap[5], and the reads subsequently split by haplotype, verifying that the copy-number alterations seen are confined to haplotype 2.*

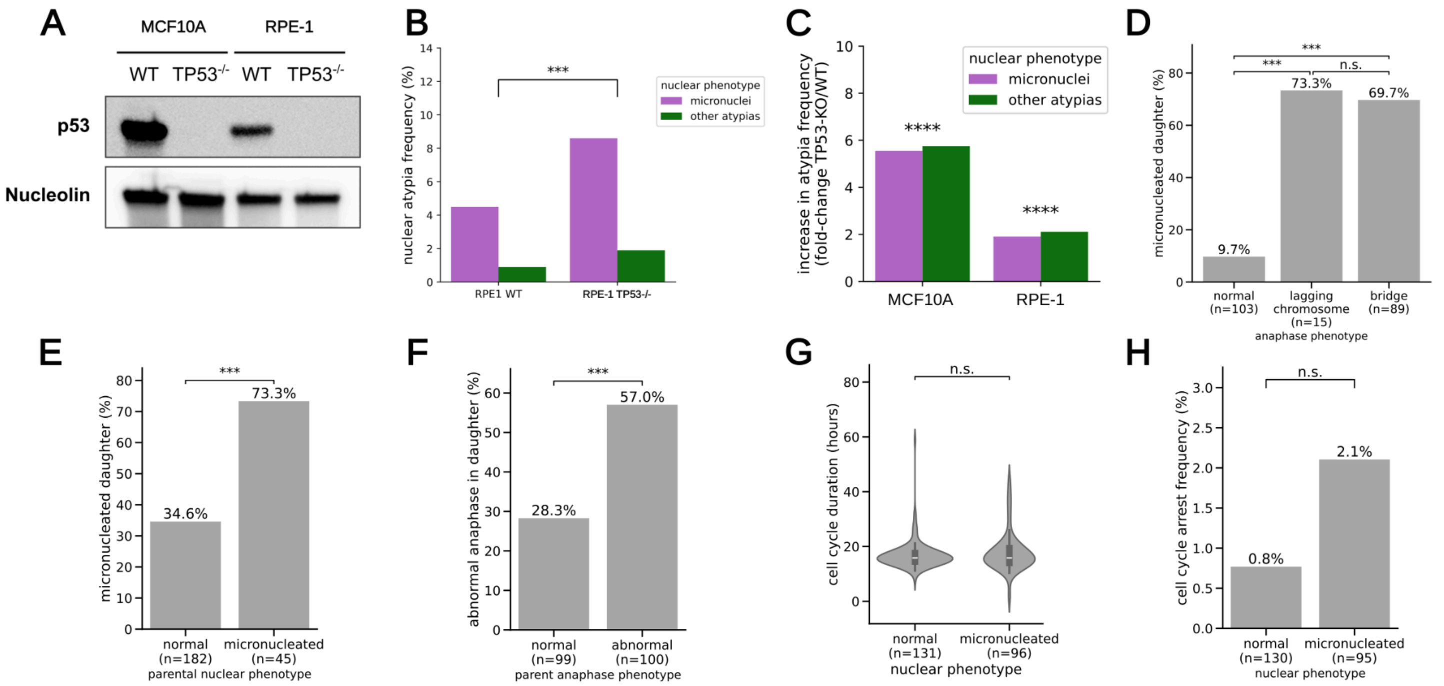


**Figure S8 | Characterisation of the *TP53-/-* cell line models used in this study**
*(A) Western blot for the p53 protein, with nucleolin used as loading control. (B) Nuclear atypia frequency in RPE-1 WT and TP53-/- cell line models (Fisher's exact test). (C) Fold-increase in nuclear atypia frequency of TP53-/- models compared to their WT counterparts (Fisher's exact test, Bonferroni corrected, P<0.0001). (D) Frequency of daughter micronucleation associated with different anaphase phenotypes (Fisher's exact test). (E) Frequency of micronucleated parent cells that generate at least one micronucleated daughter (Fisher's exact test). (F) Frequency of abnormal anaphases in daughter cells depending on the anaphase phenotype of the parent cell (Fisher's exact test) (G) cell cycle duration in hours for daughter cell generation. Duration measured from mitosis to mitosis (Mann-Whitney U test, two-sided, n.s. not significant) Center line, median; inner box limits, upper and lower quartiles; whiskers, 1.5x interquartile range; gray patch, kernel density estimate. (H) Frequency of cell cycle arrest, defined as cells with a cell cycle duration longer than the 99th percentile (Fisher's exact test).*

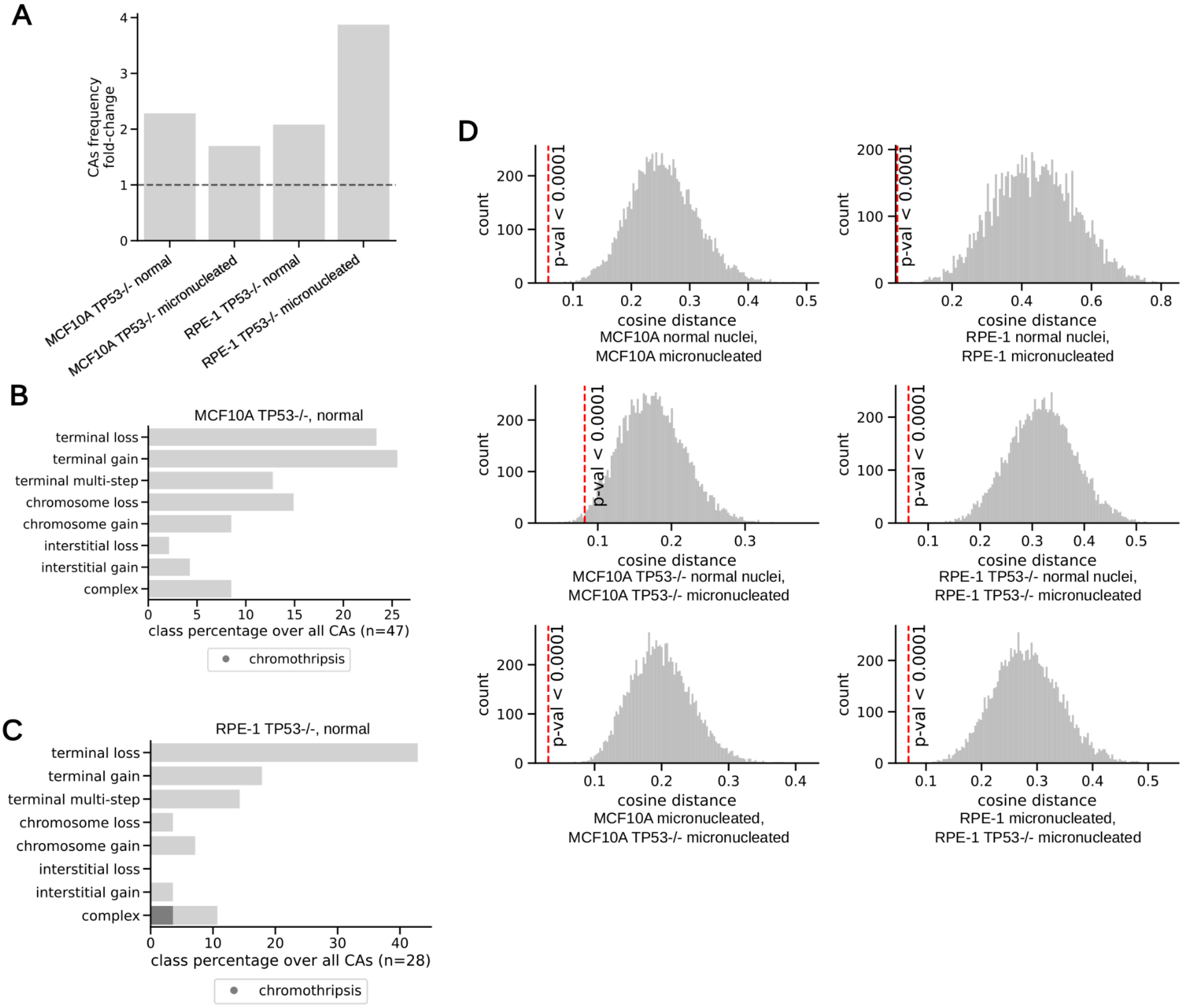


**Figure S9 | CA landscape characterization in the *TP53-/-* cell line models**

*(A) Fold-change in determined CA numbers per cell for TP53-/- versus the respectively matched WT cell line model, shown across cell line models and nuclear phenotypes. (B,C) Breakdown of CA classes in normal (i.e. not micronucleated) MCF10A (B) and RPE-1 (C) TP53-/- cells, with the class percentage shown in comparison to all CAs detected. (D) Distribution derived from permutation of CA class labels in sample pairs, with the cosine distance statistic (see* ***Methods****) used to evaluate CA class distribution similarity. Gray histogram: expected distribution of cosine distance between the reference sample and the permuted one. Red dashed line: observed statistic. The observed cosine distance is significantly smaller than expected under the permutation, implying high similarity of CA class distributions across experimental conditions.*

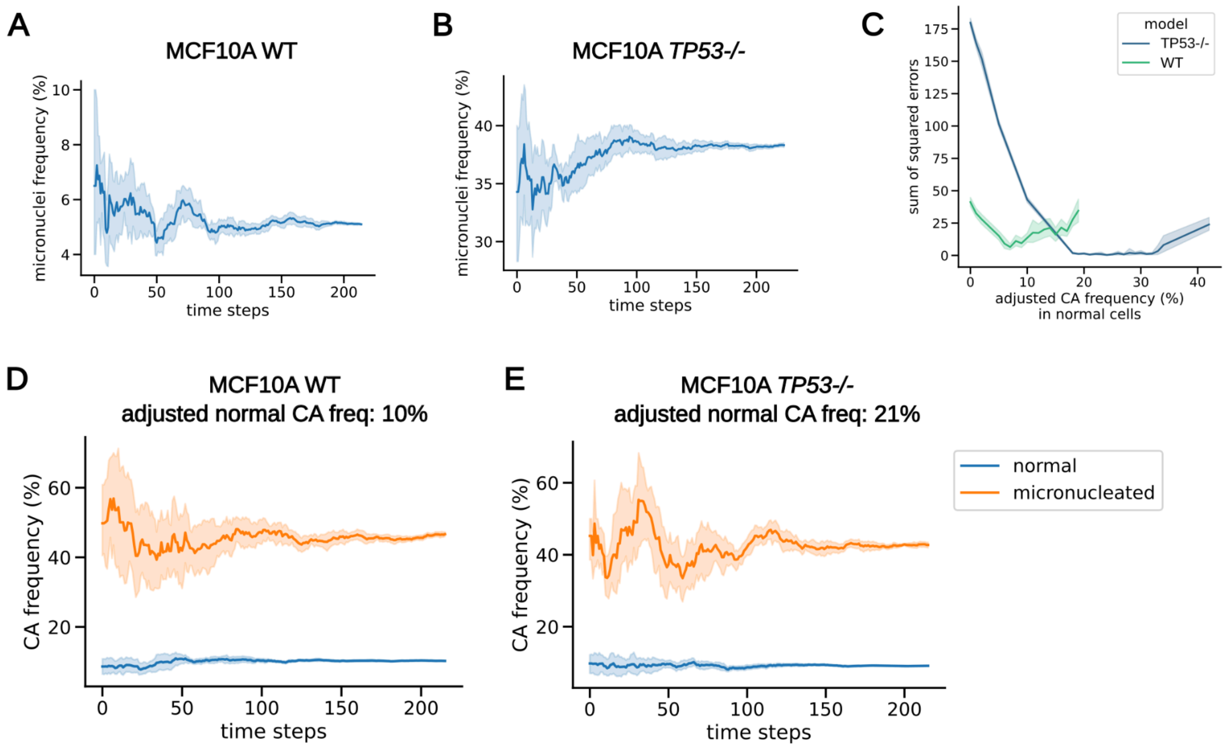


**Figure S10 | Characterization of the agent-based model for CA rate estimation**
*(A,B) Simulated micronuclei frequency for the MCF10A WT (A) and TP53-/- (B) models. Coloured band: 95% confidence interval calculated over 10 simulations. (C) Goodness-of-fit optimization for MCF10A WT and TP53-/- models with different adjusted levels of CA frequency. For each frequency level tested, optimisation was run starting from 50 initialisations using uniform random numbers. (D,E) Simulated CA frequency for MCF10A WT (A) and TP53-/- (B) models. Coloured bands: 95% confidence interval calculated over 10 simulations.*

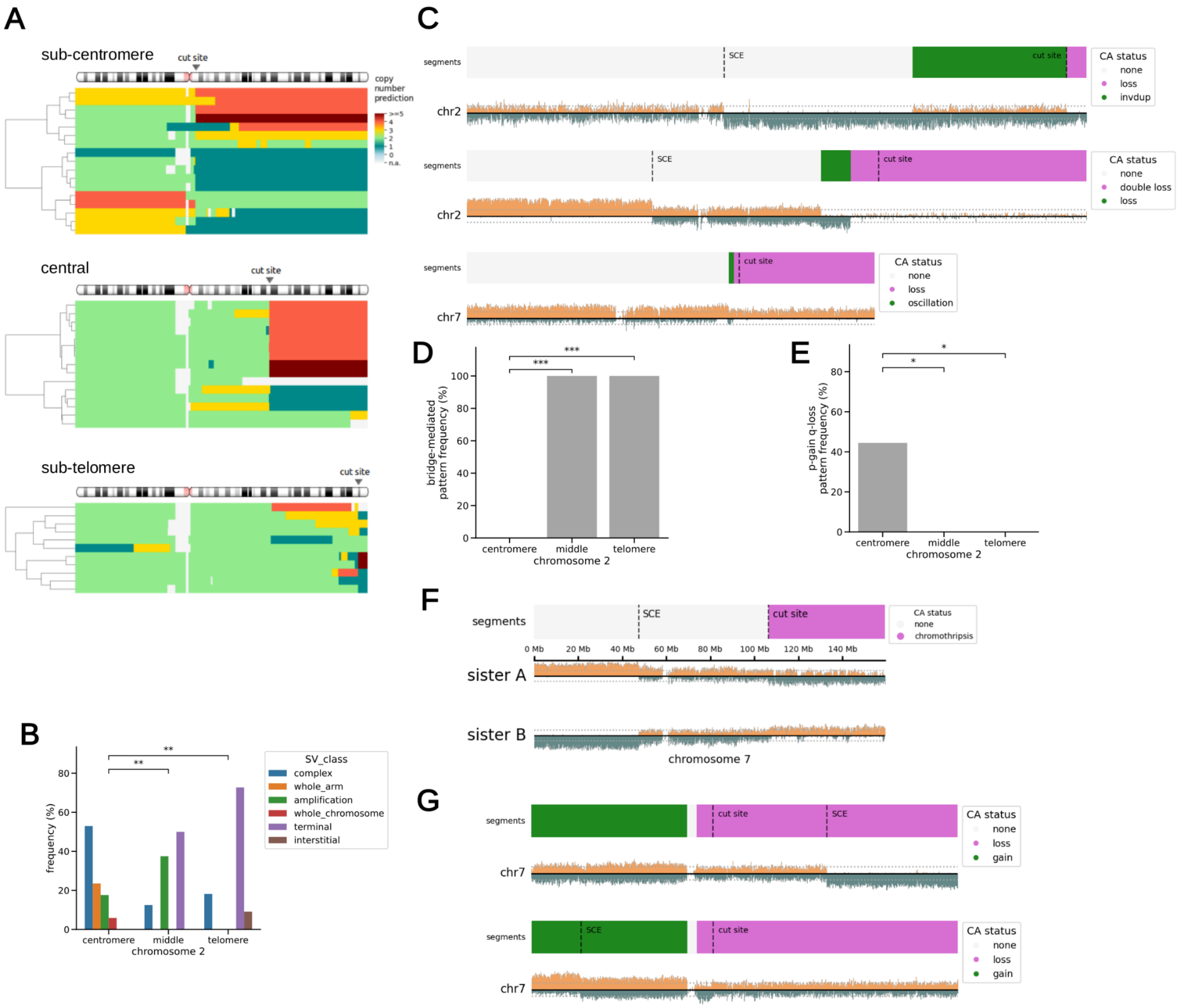


**Figure S11 | Detailed characterisation of CAs arising from targeted DSB induction**

*(A) Overview of copy-number changes for chromosome 2 q-arm CAs, shown for sub-centromere (top), central (middle) and sub-telomere (bottom) target loci. (B) Types of CA observed for different cut sites on the chromosome 2 q-arm. (C) Terminal multi-step CAs on targeted chromosomal arms. (D,E) Significant enrichment for bridge-mediated (D) and isochromosome-like (E) CA patterns, depending on the cut location. We considered all terminal and complex CAs in this analysis (P-values are based on Fisher's exact test). (F) Chromothripsis in sister cells, involving targeted DSBs at the central cut site. (G) Examples of inferred isochromosomes. Dashed lines correspond to SCEs and targeted cut sites. The chromosome 7 consensus copy number of the MCF10 cell line is three.*

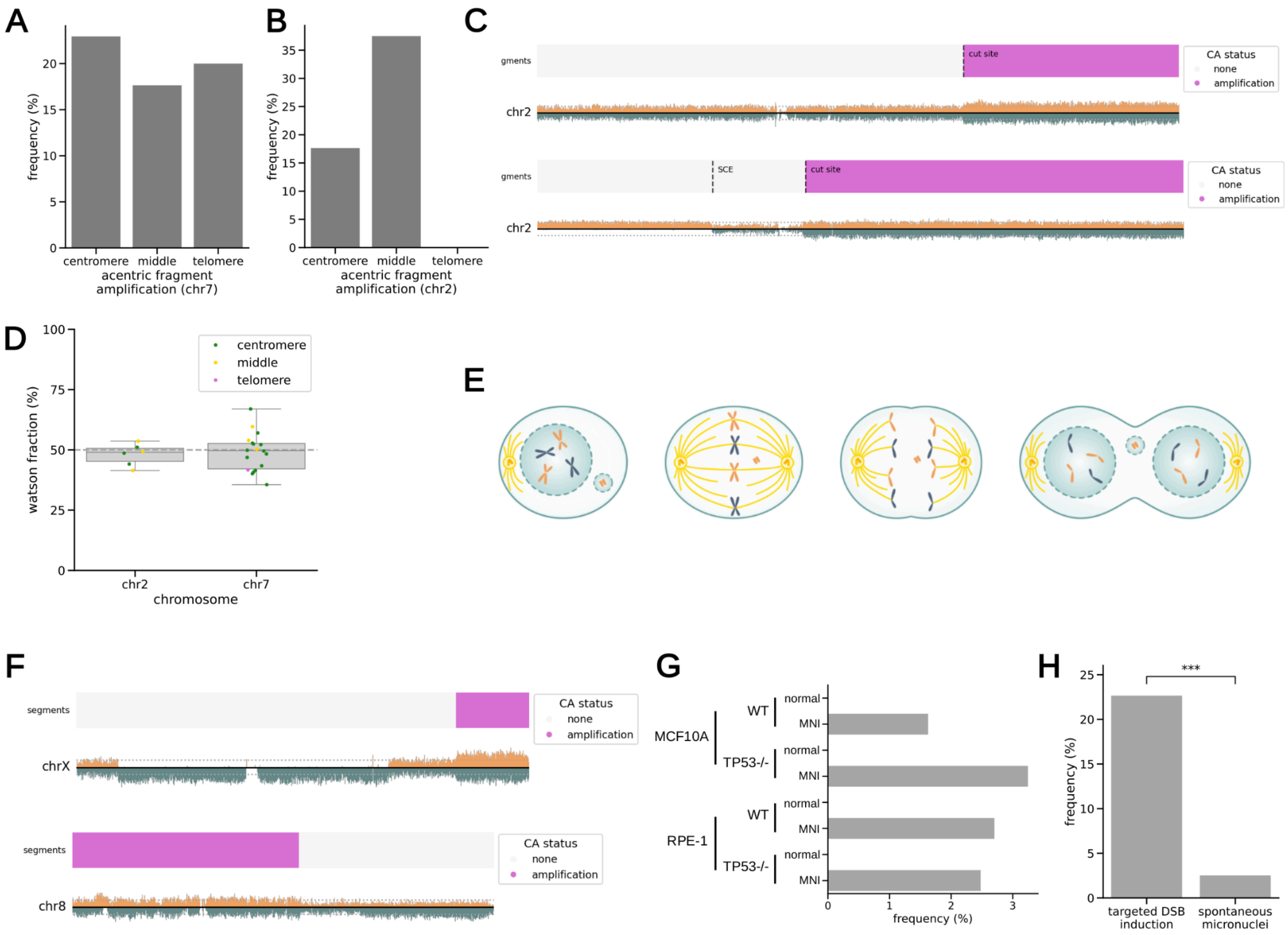


**Figure S12 | Analysis of acentric fragment amplifications**

*(A,B) Frequency of observed acentric fragment amplifications in our targeted DSB experiments for chromosome 7 (A) and chromosome 2 (B). (C,F) examples of acentric fragment amplification in spontaneously formed micronuclei (F) and following targeted DSB induction (C). (D) Watson (W) read fraction of amplified acentric fragments. (E) Scheme showing how acentric fragments contained in micronuclei do not participate in mitosis and can be inherited as a single unit. Center line, median; box limits, upper and lower quartiles; whiskers, 1.5x interquartile range; points, outliers. (G) frequency of acentric amplification in spontaneous micronuclei across cell lines and nuclear phenotypes. MNI: micronuclei. (H) Frequency of acentric amplification in spontaneous micronuclei when compared to targeted DSB induction (significance testing based on Fisher's exact test).*

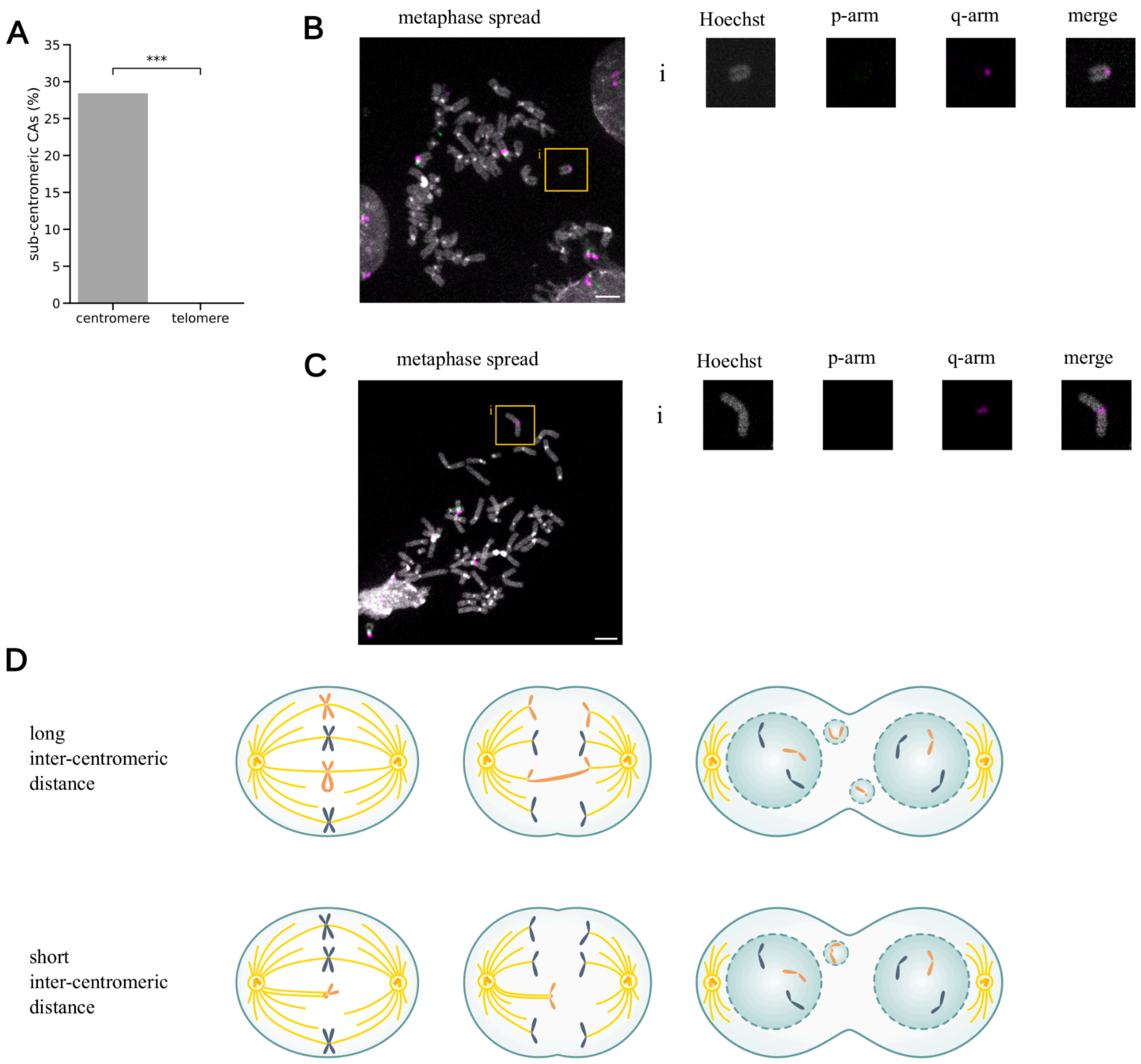


**Figure S13 | Mechanism of isochromosome generation by U-type exchange**
*(A) Frequency of abnormal sub-centromeric probe hybridisation patterns across different cut-sites (Fisher's exact test). (B) Metaphase spread example with abnormal chromosome 7 structures. Regions of interest (ROIs) are – A: cut fragment constituting the q-arms of chromosome 7 and a probe signal at one extremity. (C) Metaphase spread example with abnormal chromosome 7 structures. ROI A: isoacentric chromosome, with similar sized segments around the middle q-arm signal. (D) Mitosis schemes with isodicentric chromosomes, contrasting mitosis with chromatin bridge formation from long-intercentromeric distance (top) and isochromosome formation with short-intercentromeric distance (bottom). We note that isodicentric chromosomes might still form micronuclei following their formation, due to potential occasional erroneous kinetochore-microtubule attachments. One possibility is that isodicentrics might form monotelic spindle attachments, leading to misaligned chromosomes that satisfy the spindle assembly checkpoint and micronucleation[6].*

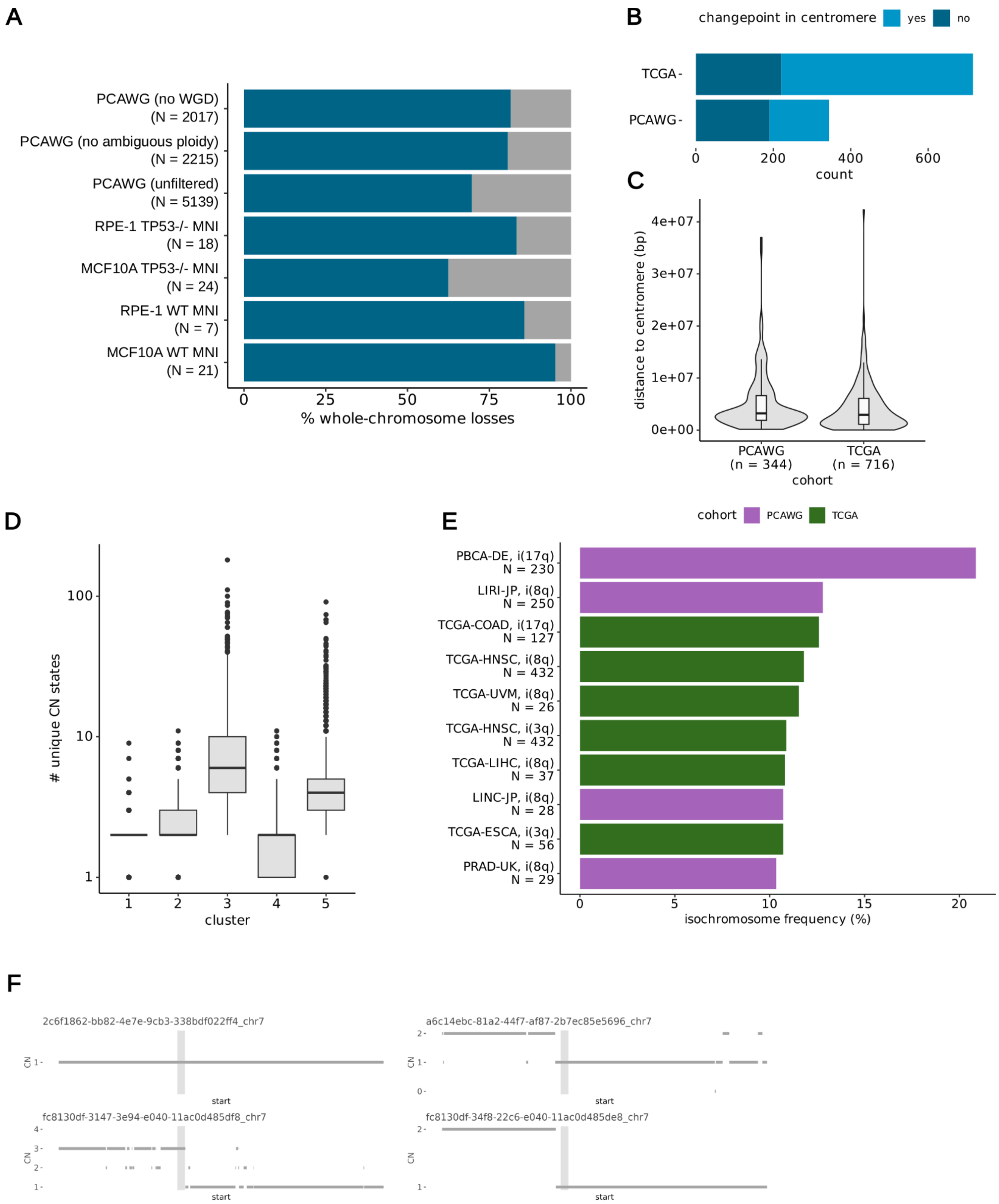


**Figure S14 | Aneuploidy and isochromosome analysis in primary cancer genome datasets**

*(A) Proportion of whole-chromosome losses among all whole-chromosome aneuploidy events in spontaneous micronuclei models and the PCAWG[7] dataset. The "unfiltered" PCAWG dataset includes all representative aliquots[7]. The "no ambiguous ploidy" dataset only includes those aliquots in which an integer ploidy could be assigned unambiguously (see* ***Supplementary Notes****). The "no WGD" dataset excludes aliquots affected by whole genome duplication (WGD). The total number of whole-chromosome gains and losses is shown for each dataset. MNI: micronuclei; TP53KO: knockout of TP53 gene; WT: wild type, for normal TP53 status. (B,C) Number of inferred isochromosomes (B)*

*and distribution of distances between isochromosome changepoints and centromeres (C) identified in the TCGA and PCAWG datasets, considering only non-redundant donors. Center line, median; inner box limits, upper and lower quartiles; whiskers, 1.5x interquartile range; gray patch, kernel density estimate. (D) Number of unique CN states per chromosome in the PCAWG dataset. Chromosomes are grouped by clusters as defined in* ***Figure S15****. Center line, median; box limits, upper and lower quartiles; whiskers, 1.5x interquartile range; points, outliers. (E) The top 10 most frequent isochromosomes identified in TCGA and PCAWG sub-cohorts with more than 25 donors. PBCA: paediatric brain cancer; LIRI: hepatocellular carcinoma; COAD: colon adenocarcinoma; HNSC: head and neck squamous cell carcinoma; UVM: uveal melanoma; LIHC / LINC: liver hepatocellular carcinoma; ESCA: Esophageal Carcinoma; PRAD: prostate adenocarcinoma. (F) Examples of terminal losses affecting the q-arm of chromosome 7 in breast cancer samples from the PCAWG cohort.*

---

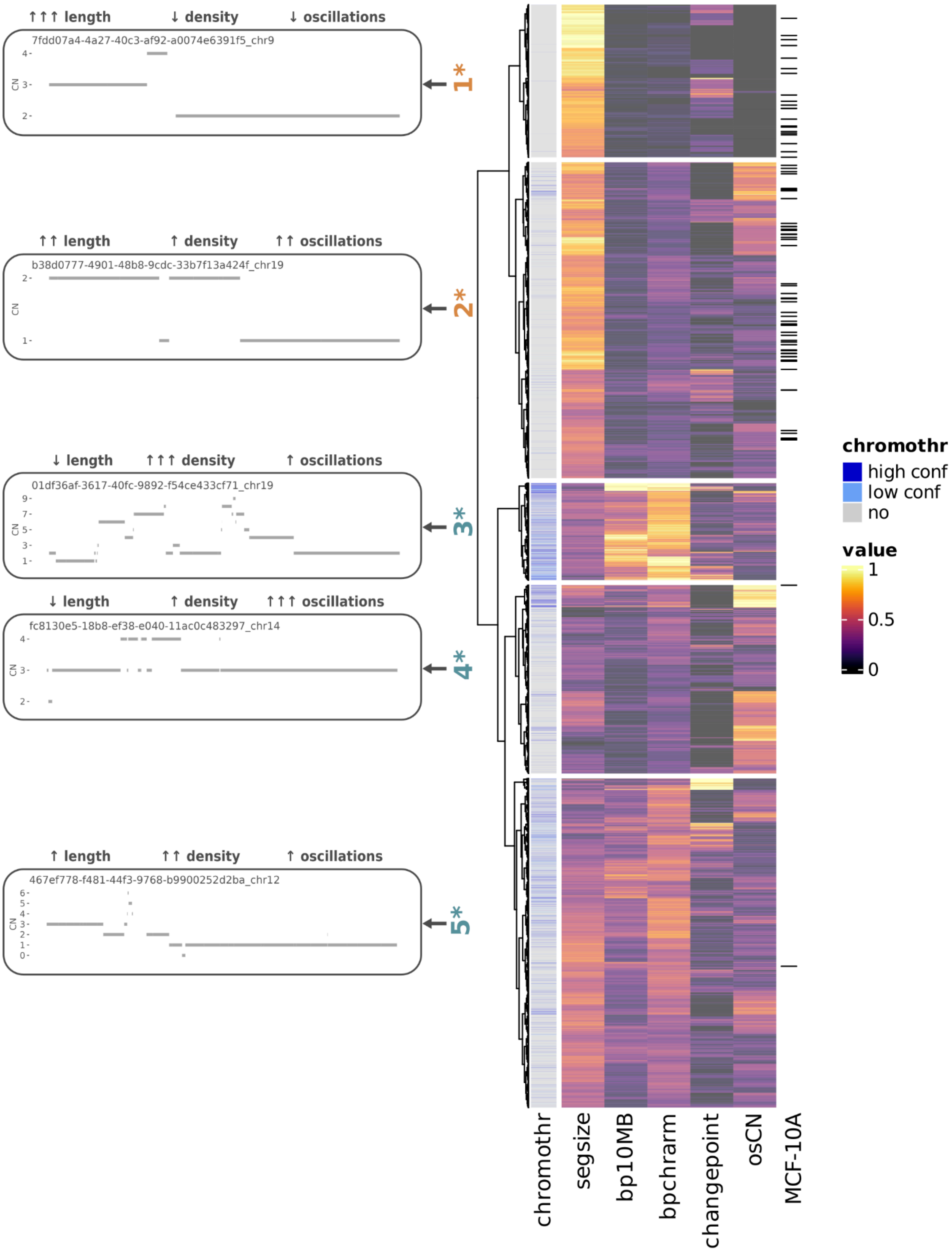


**Figure S15 | Copy-number alterations clustering based on copy-number (CN) features**
*Heatmap of CN features across PCAWG and MCF10A chromosomes with at least 2 non-diploid segments. MCF10A chromosomes are highlighted with ticks on the right side. Chromosomes are clustered by Euclidean distances and ward.D2 linkage. For visualisation purposes, 99% winsorization was applied prior to normalisation. Inferred chromothripsis events are shown on the left column bar. For PCAWG, high as well as low confidence chromothripsis calls were considered[8]. For MCF10A, we considered all of our manually curated calls (we note that owing to the single-cell nature of the MCF10A data with much fewer reads generated per genome, short chromothripsis-derived DNA fragments <200kb in size are likely to remain undetected by Strand-seq). For each cluster number,*

*asterisks denote significant enrichment (orange) or depletion (turquoise) of MCF10A chromosomes in each cluster, as assessed by a permutation test (FDR-adjusted $p < 0.05$). To the left of each cluster, a schematic of the CN patterns of a representative chromosome is shown. Chromothr, chromothripsis; segsize, length of non-diploid segments; bp10MB, number of breakpoints per 10 Mb; bpchrarm, number of breakpoints per chromosome arm; changepoint, magnitude of the changepoints, osCN, number of CN oscillations.*

---

### Supplementary Tables

**Supplementary Table 1.** Summary of single-cell clones analysis by low-pass WGS, with CA annotations. (Table accompanying the manuscript as spreadsheet)

**Supplementary Table 2.** Summary of sgRNA sequences and PCR primers for the TIDE assay. (Table accompanying the manuscript as spreadsheet)

**Supplementary Table 3.** Convolutional neural network architecture (Table accompanying the manuscript as spreadsheet)

**Supplementary Table 4.** Data sources for PCAWG and TCGA analyses. (Table accompanying the manuscript as spreadsheet)

**Supplementary Table 5.** PCAWG and TCGA cohort identifiers data. (Table accompanying the manuscript as spreadsheet)

**Supplementary Table 6.** Isochromosome annotation and results. (Table accompanying the manuscript as spreadsheet)

**Supplementary Table 7.** CA annotations for the targeted DSB-induction MAGIC experiments at the *HPRT1* locus on chromosome X. (Table accompanying the manuscript as spreadsheet)

**Supplementary Table 8.** CA annotations for the MAGIC experiments with the MCF10A and RPE-1 WT and *TP53-/-* cell line models, for micronucleated and normal cells. (Table accompanying the manuscript as spreadsheet)

**Supplementary Table 9.** CA annotations for targeted DSB-induction MAGIC experiments along chromosome arms 7q and 2q. (Table accompanying the manuscript as spreadsheet)

### Supplementary Methods

#### Photolabeling strategies

In this study, we employed two photolabeling strategies based on Dendra2 and DACT-1. Dendra2 is a monomeric fluorescent protein, originally derived from *Dendronephthya sp*., that is amenable to photoconversion with purple-blue light[9]. The photoconversion reaction is irreversible and thus duration of the labeling depends on protein turnover. In our case, we chose to employ H2B-Dendra2, where the photolabeling protein is fused to the histone 2B protein[10]. This enables the visualisation of nuclear atypias, as well as photoconversion. Moreover, thanks to the low turn-over of histone proteins, photolabeling persists also across subsequent generations as demonstrated by the collection and single-cell analysis of sister cells. DACT-1 is a dual-activatable cell tracking molecule previously developed for the controlled and prolonged tagging of single-cells[11]. DACT-1 is cell permeable and accumulates inside the cytoplasm thanks to the action of an esterase that converts it into its pre-activated form. The dye is dark in its native state, but a photolysis reaction induced by 405 nm light causes the conversion of pre-activated DACT-1 into a bright red-fluorescent molecule, which also forms a cross-link with intracellular nucleophiles. These features ensure that DACT-1 can only be fully activated in the intracellular space, with cross-linking extending the stability of photolabeling. We synthesised DACT-1 in house, in EMBL's Chemical Synthesis Core Facility. In the case of one experiment with RPE-1 cells, we used DACT-1 as photolabel. In this case, nuclei were visualised with NucSpot Live 650 Nuclear Stain (Biotium). Both molecules were added to the cell culture medium at least three hours prior to the start of the photolabeling phase. Photolabeling for both H2B-Dendra2 and DACT-1 was performed by line scanning inside regions of interest covering the parental nucleus, with a 20X objective, 405 nm laser at 0.5% power, in the low intensity range, for 50 iterations.

#### Microscope automation and imaging

MAGIC relies on full microscope automation and computer vision, to enable large-scale, phenotype-driven targeted illumination of single-cells based on its adaptive-feedback loop. Adaptive-feedback microscopy can be understood as sequences of imaging and analysis jobs, where each step in the sequence depends on the execution and actionable information coming from the previous steps. This requires the controlled interaction between microscope hardware and computer vision algorithms to generate complex behaviours required for a fully automated experiment[12]. For this purpose we developed three main software components: the first component is a microscope control script that controls the automation workflow, integrates feedback information and directly controls the microscope hardware. The second component, based on *AutoMicTools*[13], manages image analysis tasks by interacting with the microscope control component, performing the analysis tasks and generating feedback files. The third component is a python package called magic_tools, which we developed to extend the image analysis capabilities of our platform with ML; magic_tools offers an image processing server, which runs on a cluster node and takes care of computationally intensive tasks such as the identification of micronuclei.

For our MAGIC experiments we employed an LSM 900 microscope (Zeiss) equipped for confocal and widefield imaging. Microscope automation was implemented requiring two software components: a custom script for controlling the microscope and an ImageJ plugin to handle the communication between microscope and image analysis server. Control of the microscope was accomplished through the developer toolkit of the software ZEN, enabling control via scripts written in IronPython. We designed a custom script that implements the feedback loop as described in **Figure S2A**, triggering imaging experiments and coordinating between imaging and analysis operations. At the same time, image analysis tasks are accomplished via *AutoMicTools*[13], a Java library that works as an ImageJ plugin and coordinates with the microscope control script. For the purpose of this project, we additionally developed an *AutoMicTools* Job class that handles image analysis via the *magic_tools* image processing server. The microscope control software and *AutoMicTools* interact by exchanging files in a “watchfolder” where images and json files are monitored to trigger acquisition or analysis jobs. Both custom software components are made available in the magic_automation repository.

The microscope control script iterates through several positions to perform three tasks: autofocusing, identifying micronuclei, and photoconverting target nuclei. Autofocus is accomplished by finding the reflection of the glass bottom dish. To achieve this, the microscope acquires an XZ line scan with a 639nm laser. The image is then detected and analysed by *AutoMicTools* to look for the location of highest intensity along the Z axis, representing the reflection of the glass. A feedback file is generated containing the coordinate of the best-focused slice and this information is integrated in the next task. To identify micronuclei, we use a 20X/0.8 NA M27 air objective and a 488 nm laser to acquire a Z-stack image. This image consists of seven equidistant slices spanning a total of nine µm, centred on the focus coordinate identified in the previous step. *AutoMicTools* detects the image and sends it for analysis to an image analysis server based on magic_tools. The classification result is converted in polygon coordinates that define the contour of the nuclei to be photolabeled. Finally, for targeted photolabeling, the script parses the feedback file containing the polygon coordinates and defines regions of interest (ROIs) as in a bleaching experiment, before the experiment is triggered. Photolabeling is performed with a 405 nm laser scanning through the ROIs, for 50 iterations each, at speed 8 and 0.5% power. Two images before and after the experiment are acquired. At this point the microscope proceeds to the next position for a new iteration.

During a typical experiment we define 200 fields of view per well, for a total of 800 positions, which are scanned over the course of 24 hours or until the desired yield is reached.

#### Online image analysis with *magic_tools*

MAGIC necessitates computer vision inferences to achieve online, real-time image processing. Yet, the calculation of pixel and object-level features for phenotype classification is a computationally-intensive task that scales linearly with the number of pixels and objects to be evaluated as well as the number of features. Therefore, we developed *magic_tools* to enable parallelization of feature computation by means of an image processing server, and to support annotations and training of ML models.

For real-time detection of micronucleated cells, we integrated an image analysis pipeline into magic_tools, which includes a two-step semantic pixel segmentation and object classification based on extreme gradient boosting classifiers[14] trained on precomputed mathematical features. In the first step, we perform a maximum-intensity projection of the Z-stack and use an XGBoost classifier to perform a pixel semantic segmentation. For semantic segmentation, we compute pixel-level features by applying a series of Gaussian filters to capture information at different scales, and then filters for pixel intensity, edgeness and texture to generate the final feature outputs. An XGBoost classifier then generates a foreground prediction score for each pixel. Semantic segmentation is refined by morphological operations and on-demand watershed[15] to define nuclei objects. At this stage, micronuclei – if separated from the main nucleus – might form their own object or be part of a nuclear mask if they are adjacent to the nucleus. Therefore, we refine the nuclei masks to identify areas of convexity that deviate from the oval shape of a normal nucleus and thus capture micronuclei-candidate objects. We calculate object-level features for micronuclei candidates across multiple domains: basic descriptors, shape, intensity and texture. In addition, we calculate the same features for parental nuclei. Micronuclei and parent nuclei features are then concatenated by nearest-neighbour search, considering the shortest distance between a micronucleus and a nucleus edge. The concatenated feature vectors are then used as input for the second XGBoost classifier to identify micronuclei among all the candidate objects. Objects that cross a predefined probability threshold are considered eligible for targeted illumination and the parental nuclear masks are converted to polygon coordinates and packaged in a feedback file.

For parallelisation of feature computations, *magic_tools* uses two different approaches in the case of pixel or object-level features. In the case of pixel features, filter computations are parallelised on a pool of workers in two stages, first across Gaussians, then across feature filters. Instead, in the case of object features, parallelisation occurs across objects, with all features calculated in sequence for each of them. This parallelisation enables inferences in the order of a few seconds, depending on the number of objects in the image. Another feature facilitating real-time inference is the image processing server, which we deployed on a computational support node to leverage more computational power. The image processing server can receive images via a RESTful API and we developed a custom *AutoMicTools* plugin to enable online image analysis by *magic_tools*.

Image analysis for round nuclei detection utilises the same approach, but it uses a round-nucleus specific XGBoost classifier. In alternative, we selected cells with a low probability of carrying a micronucleus. All image analysis pipelines are available in the *magic_tools* repository.

#### Strandtools – an optimised single-cell CA calling algorithm for Strand-seq data

To detect CA in single-cell genomic data generated by Strand-seq, we developed a python package called *strandtools* (https://git.embl.de/cosenza/strandtools), that improves on our previously developed tool MosaiCatcher[1], and is tailored for the specific task of handling CA detection in single cells and complex ploidy backgrounds. In Strand-seq data, the ratio in read counts in the W and C strand orientation states is related to the underlying number of template homologs, with specific ratios providing information on segmental copy numbers and SV classes based on the principle of tri-channel processing (combining information on read depth, strand-orientation and haplotype phase)[1]. Particularly, strandtools leverages read coverage, as well as the ratio between W and C templates from multiple cells, to build Naive Bayes ML models and assign copy-number states in each single-cell, including in scenarios with a non-disomic ploidy.

While an overview of the method is provided below, we anticipate a future publication elsewhere covering in detail all of the various features of *strandtools*. In particular, data processing uses read count tables generated by MosaiCatcher with a multi-step normalisation being performed as the first step[16]. The chromosomes are then segmented in regions of homogeneous coverage and strand orientation with a modified circular binary segmentation[17] algorithm. An OPTICS clustering algorithm[18] is then employed to identify clusters of homogeneous read count and W fraction, corresponding to combinations of templates and copy number. Cluster labels are used as ground truth to train a naive Bayes classifier and generate a representation for copy number assignment when the template ratio is informative. Then, the same genomic segments have the template ratio masked to train a second naive Bayes classifier to generate copy number predictions for segments with homogeneous strand states (only W or only C reads). Copy number assignment to the different clusters is solved by brute force, maximising the correlation between copy number assigned to the different clusters and their average read count. These two Naive Bayes classifiers are then applied to the entire data to generate copy number predictions. Adjacent genomic segments with identical copy number are iteratively merged into super-segments, which can be used for a second round of classifier training for better accuracy. Several quality control (QC) filters are applied to the predictions to generate the final copy-number probabilities: we exclude segments with uncertain predictions, a too high coefficient of variation as a measure of noise, or a too high number of low count chromosomal bins. Segments that are part of excluded genomic regions (e.g. centromeres) are likewise excluded. Finally, changepoints for copy-number breakpoints and SCEs are outputted. In our workflow, this output is used for detailed CA classification as described in the **Methods** section. The source code of *strandtools* is provided alongside this study (see Code availability section).

#### Computational sister cell pair discovery from Strand-seq data

To automatically detect sister cells, *strandtools* converts the strand state class into a numerical value: 1 for WW, 0 for WC and -1 for CC status. Then, the Pearson correlation coefficient between all pairs of cells in an experiment is computed. Sister cells are considered as cell pairs exhibiting a correlation coefficient below -0.9, which we find faithfully corresponds to sister cell pairs (**Fig. S3F, S4A**).

#### XGBoost classifiers training

XGBoost classifiers are employed in two crucial moments of the analysis pipeline: pixel semantic segmentation to detect foreground signal, and object classification to identify micronuclei and their parental nuclei. For semantic pixel segmentation, we performed a sparse annotation for foreground and background pixels in seven representative images, acquired with identical imaging settings used in all MAGIC experiments with a specific cell line. We then computed a set of pixel level features, as described above, to generate our ground truth set. We split the set for training (80%), validation (10%) and testing (10%) purposes. An XGBoost classifier was trained and hyperparameters were iteratively adjusted on the validation set. Finally, we assessed classifier performance on the test set.

For micronuclei classification in MCF10A, we manually annotated 200 random fields of view and integrated this with micronuclei candidate regions detected by image analysis. This resulted in the identification of more than 23,000 candidate micronuclei regions, of which 478 were ground truth micronuclei annotations. Object level features were calculated as described above (a complete list of features can be found in the repository). Also in this case, we split the set for training (80%), validation (10%) and testing (10%) purposes, stratifying the splits by ground truth label. Model hyperparameters were fine-tuned on the validation set, to achieve at least 90% precision, while still recalling at minimum 50% of all micronuclei, striking an acceptable balance between sensitivity and specificity. Final classifier performance was assessed on the test set. To get a more objective measure of classifier performance, we performed several different dataset splits. The average precision-recall curves, with their standard deviation, were calculated from the aggregated data and presented in **Figure S2H**. We developed pixel and micronuclei classifiers tailored for the specific cell line models and to accommodate the different image settings, cell morphologies and experimental variation. Models are made available in the *magic_tools* repository.

#### Convolutional neural network based micronucleus classifier training

We additionally implemented a convolutional neural network for micronucleus classification, with the network architecture shown in **Supplementary Table 3**. The training dataset consisted of fluorescence microscopy images cropped to 300×300 pixels. The image crops were centred either on a micronucleus or a non-micronucleus object and were generated with the image analysis pipeline for micronuclei detection described above. During training, image crops were augmented using random upsampling by a factor from 1.1 to 2.0, as well as rotation and random flipping. Images were rescaled to 128×128 prior to being utilised as an input to the network. We trained the network with a negative log likelihood loss function on the image class ($L_{NLL}$) and a negative log likelihood loss between the outputs of the network for two different data augmentations ($L_{consistency}$) to encourage rotation and scale invariance. The final weighted loss function of the network was $L = L_{NLL} + 0.1 \cdot L_{consistency}$. We trained the network with 5 independent initialization and train-test splits, with 15% of the data per class (micronucleus; no micronucleus) used to construct a withheld test set. Models and training scripts were implemented using Python 3.10, jax version 0.3.6 [19] and Pytorch version 1.11.0 [20].

#### Designing single-guide RNA

Non-repetitive regions of 2–6 kb in size were identified within the centromeric and telomeric regions of chromosome 2q, which has a relatively low repeat content[21], as well as for chromosome 7q. For each site, we designed at least 10 single-guide RNAs (sgRNA) targeting these non-repetitive regions using the CRISPick tool (https://portals.broadinstitute.org/gppx/crispick/public), designing three sgRNAs per genomic region. On-target and off-target effect of sgRNAs were assessed by CRISPR-CUSTOM (https://eu.idtdna.com/site/order/designtool/index/CRISPR_CUSTOM) and Off-Spotter (https://cm.jefferson.edu/Off-Spotter/) tools. We aimed at an on-/off-target score of ≥50 (CRISPR-CUSTOM) and no binding per one, two, and three mismatches (Off-Spotter). The editing efficiency of the designed guides was assessed in sequence, using the TIDE assay[22]. For this purpose, two primers per sgRNA were designed by Primer-BLAST such that forward primer binds 150-300 bp upstream of Cas9 cut site and that reverse primer binds 300-500 bp downstream of Cas9 cut site. Designed guides showed generally good efficiency. In any case, sgRNAs showing highest editing efficiency and corresponding PCR primers, which produced a single PCR product as well as high Sanger-seq quality, were chosen for downstream analyses.

#### Statistical testing for biases in CA frequencies, SCEs and breakpoint locations

*Binomial testing for chromosome bias in CA landscapes*. To determine non-random bias in chromosomes affected by CAs, we first derived the expected probability of having any chromosome hit by a CA by dividing the number of annotated CAs and the total number of chromosomes in MCF10A or RPE-1 cells. From this, under the assumption that any chromosome would be equally affected by a CA, we derived a binomial distribution for the different chromosomes depending on their consensus copy number. We executed binomial tests for each chromosome, considering the observed CA count as the number of successful trials. Finally, P-values were adjusted by the Benjamini/Hochberg method.

*Permutation testing*. Breakpoint locations of SCEs and copy-number changes were detected using *strandtools*. To test for breakpoint enrichment in fragile sites, early and late replicating regions, we performed a permutation test utilising the regioneR package available in Bioconductor. Breakpoint regions were permuted with the randomizeRegions function 10,000 times. As evaluation functions, we employed the number of overlaps (numOverlaps) for fragile sites, early and late replicating regions, while for G4 quadruplexes we summed their occurrence in the permuted bins affected by a breakpoint. P-values were adjusted by the Benjamini/Hochberg method. Fragile site locations were obtained from HumCFS database[23], and G4 quadruplex data was obtained from a previous study[24]. We find no CA enrichment for BrdU fragile sites, consistent with prior works reporting that the low concentration BrdU pulses used in Strand-seq do not enhance genetic instability[25].

To determine enrichment for early and late replicating regions, we employed the pre-processed UW Repli-seq tracks available on the USCS Genome Browser database[26]. The data was remapped to the hg38 genome using the UCSC LiftOver tool, binned in 200 kbps windows and aggregated over the different cell lines. We calculated the mean replication timing score and its standard deviation for all genomic bins. In order to filter regions replicating early or late across cell lines, we set a low standard deviation (score standard deviation =< 5) as requirement, and then assigned bins to replicating early or late replicating regions depending on their average score (early =< 20, late >=60). Permutation tests for chromosome size took into account consensus copy numbers for each chromosome. CA annotations – regardless of the specific CA class – were permuted over all chromosomes in the dataset. For the test statistic, CA counts per chromosome were aggregated and correlation to chromosome length was computed using the Pearson correlation coefficient. P-values were then calculated from our observations, and the permuted distribution derived from 50,000 iterations.

*Binomial testing for coordinated Watson(W)/Crick(C) gains.* To test for the coordinated amplification of W and C templates, we focused on gains with an increment of exactly two. If the two extra segments were inherited independently, then their expected probability in a daughter cell would be 0.25 W/W, 0.50 W/C, and 0.25 C/C, respectively[1]. By considering every cell division as its own Bernoulli trial, and considering the W/C outcome as 'success', we employed a binomial test to examine if the observed data agrees with this model. In this case the probability of success for a trial is 0.5, the number of trials corresponds to the number of cells carrying an amplification, and the number of successes is the number of cases where we observe a WC ‘coordinated gain’. Among 18 inferred acentric gains with a copy-number increment of 2, the W/C ratio remains 1:1 in all 18 cases, based on which we reject the hypothesis that the fragment gains segregate independently ($P$<7.63e-06). We thus infer based on the single–cell genomic data analysed that the amplified segments are part of the same derivative chromosome, where they are arranged in an inverted orientation.

### CA rate estimation by bound-constrained minimization

To estimate the CA rates $R$, we performed a bound-constrained minimization of the sum of squared errors between simulated and target *de novo* CA frequencies. To achieve this, we employed a modification of the “Powell” method[27]. This optimization algorithm searches the parameter space by doing a sequential, bi-directional minimization for each search vector, until a local minimum of the function is found.  As rates $R$ are probabilities of an event happening, we bound these values between 0 and 1. For each estimation, we ran 50 optimization rounds, initialising the $R$ parameters from the same set of uniformly random numbers. The output rates of one estimation were computed by the average of the 50 optimization outputs, weighed by the sum of squared errors.

Under the hypothesis that not all observed CA would be *de novo*, we adjusted the target CA frequencies by subtracting a scalar value from the observed CA count in both normal and micronucleated cells. We performed first a sparse search, then focused on the following ranges for normal cells: 5 to 19% for WT and 18 to 34% for *TP53-/-* models. We thus derived CA rate estimates for each of the adjusted frequency values with their corresponding sum of squared errors values. We used this latter value as a measure of goodness-of-fit to judge how well the model fitted the observations, and determine the final CA rates reported in **Figure 3G**.

### Western blotting

Western blotting to validate p53 knock-out status of cell lines was performed as follows: a pellet from 2 million cells was resuspended in lysis buffer (1X RIPA buffer Cell Signaling, 1X protease inhibitor cocktail Roche, 1mM PMSF Roth) and incubated on ice for 10 minutes. Debris was pelleted and the supernatant was frozen for 2 hours at -80C to assist lysis. 40µg of protein were mixed with loading dye and heated for denaturation. Samples were loaded onto a precast 4-15% Tris-Glycine gel (Bio-Rad) and run in 1X Laemmli buffer at 100V for 10 minutes, followed by 180V for 45 minutes. We then performed semi-dry transfer onto a nitrocellulose membrane with a Trans-Blot Turbo Transfer System. After transfer, the membrane was blocked with 5% milk in TBS-T and primary antibodies (1:200 p53, DO-1 mouse mAb, Santa Cruz; 1:1000 nucleolin, D4C7O rabbit mAb, Cell Signaling) were incubated for 2 hours in blocking buffer. The membrane was washed three times in TBS-T and an HRP-coupled secondary antibody (goat anti-mouse IgG-HRP 1:10000, Invitrogen) was incubated for 45 minutes. After three final washes, the membrane was incubated in ECL substrate (Bio-Rad) for 1-3 minutes at room temperature in the dark. Protein bands were visualised using a GelDoc XR Imaging system.

### Supplementary Notes

#### Pan-cancer WGS cohorts

Data from the Pan-Cancer Analysis of Whole Genomes (PCAWG) Consortium[7], encompassing 2,583 donors across 47 tumour types, was downloaded from the International Cancer Genome Consortium (ICGC) Data Portal (https://dcc.icgc.org/releases/PCAWG, **Supplementary Table 4**). To avoid redundancies due to multiple aliquots originating from the same donor[7] we selected one representative aliquot per donor (as defined in "aliquot_donor_tumor.whitelist.tsv.gz" in the ICGC Data Portal). For the copy number (CN) data, we merged consecutive segments that had the same CN value.

To comprehensively infer isochromosomes across diverse cancer types, we combined CN data from PCAWG[7] and from a separate TCGA pan-cancer WGS resource, the latter comprising data from 4,957 tumour-normal pairs encompassing 30 tumour types (https://portal.gdc.cancer.gov/). For TCGA, WGS datasets with an average read length of ≥100 bp and passing the following quality control (QC) criteria were considered for our analyses. QC was conducted using AMBER and PURPLE, with samples discarded if at least one of the following conditions was met: (1) FAIL_CONTAMINATION, where tumour contamination in homozygous sites from the normal sample exceeded 10%; and (2) FAIL_NO_TUMOUR, where no evidence of tumour was found in the sample. For the 698 donors that were found in both the TCGA and PCAWG cohorts, only the TCGA sample was kept for isochromosome analysis (n=1,985 for PCAWG; n=4,957 for TCGA). Raw sequencing reads were mapped to the GRCh38 build of the human reference genome using BWA-MEM (v0.7.17-r1188).[28] Aligned reads were processed following the GATK Best Practices workflow (v4.1.8.0) to remove duplicates and recalibrate base quality scores[29]. Germline and somatic SNVs were called and filtered using SAGE (v2.8). Somatic SVs were detected using GRIDSS2 (v2.12.0, available at https://github.com/PapenfussLab/gridss)[30] and filtered using GRIPSS (v1.9). The B-allele frequency (BAF) of heterozygous SNPs was computed with AMBER (v3.5) and read depth ratios were calculated using COBALT (v1.11). B-allele frequency, read depth information, breakpoint coordinates and single nucleotide variant (SNV) allele frequencies were integrated to estimate somatic copy number aberrations using PURPLE (v2.54).[31] The raw copy number values estimated for each segment were rounded to their nearest integer for further analysis. Then, consecutive CN segments that had the same CN value were merged. Only copy-number values inferred by PURPLE using BAF data were considered for downstream analysis. SAGE, GRIPSS, AMBER, COBALT, PURPLE and LINX were developed by the Hartwig Medical Foundation (HMF)[31] and are freely available on GitHub at https://github.com/hartwigmedical/hmftools.

#### Comparing copy-number features from spontaneous micronuclei with the PCAWG resource

##### Extracting copy-number features for abnormal chromosomes

We compared copy-number (CN) features between MCF10A and the PCAWG WGS dataset[7] by adapting methods from Drews and colleagues[32] (*i.e.*, the CINSignatureQuantification package; https://github.com/markowetzlab/CINSignatureQuantification#cinsignaturequantification) to facilitate their application to a single-cell genomic dataset. These methods quantify five CN features: (1) the length of non-diploid segments; (2) number of breakpoints per 10 Mb; (3) number of breakpoints per arm; (4) number of CN oscillations; and (5) magnitude of the changepoints. In particular, we adapted the code to allow for an extraction of CN features at a per-chromosome level, and provided as an input CN calls (CN integer values). For MCF10A, we modified the input CN data by computing max(0, CN - consensus + 2), so that non-diploid segments whose CN was the same as the consensus were not considered to be CAs. The adaptations to CINSignatureQuantification introduced included skipping the smoothing of segments by smoothSegments() when the input data only has one segment, and setting DCIN=2 in removeQuietSamples() so that chromosomes harbouring fewer than 2 non-diploid segments are filtered out (this is the lowest possible attainable value within the framework of this available software package). In addition, for the computation of CN oscillations, we noted an unwanted behaviour in the original getOscillationDrews() function from the CINSignatureQuantification package by which oscillations were not counted if they reached the end of the chromosome. We adapted the code to prevent this behaviour. After computing chromosomal instability (CIN) features[32] using the adapted software, we averaged the log2-transformed values (adding a pseudocount of 1) for each feature in each chromosome. Finally, we normalised all values across each feature between 0 and 1. We did the normalisation separately for three datasets of interest – MCF10A, PCAWG, and combined (MCF10A + PCAWG).

We performed hierarchical clustering of the CIN features using Euclidean distances and ward.D2 linkage as implemented in the hclust() function in R. For each heatmap, we determined the optimal number of clusters using the gap statistic as implemented in cluster::clusGap() with factoextra::hcut(). Then, we plotted the heatmaps using the ComplexHeatmap package. For plotting purposes, we applied 99% winsorization to the mean log2-transformed feature values prior to normalisation. In the combined clustering of PCAWG and MCF10A chromosomes, we assessed the enrichment or depletion of each cluster in MCF10A chromosomes by permuting the cluster labels 1000 times and comparing the observed proportion of MCF10A chromosomes in each cluster with the distribution of simulated values. Finally, we corrected the P-values for multiple testing using the Benjamini-Hochberg method.

#### PCAWG based copy-number (CN) pattern cluster enrichment analysis

To assess similarities between the CA patterns in MCF10A and those observed in the primary cancer genomes[7], we utilised the methodology adapted from[32] as described above. For each chromosome that had at least two non-diploid segments, we computed five features related to non-diploid segment length, number of CN breakpoints per 10 Mb, number of CN breakpoints per arm, number of CN oscillations, and magnitude of CN changepoints. Next, we performed a joint clustering of these features computed on PCAWG and MCF10A chromosomes (**Fig. S15**). We find that MCF10A chromosomes cluster together with PCAWG chromosomes, leading to five major clusters.

Cluster 1 is characterised by relatively simple chromosomes, harbouring few breakpoints and long segments. Cluster 2 contains more complex chromosomes than cluster 1, harbouring comparatively shorter segments and higher breakpoint density, as well as medium/long chains of oscillations. Cluster 3 contains highly complex chromosomes, harbouring short non-diploid segments, the highest breakpoint density, and the highest magnitude of CN changepoints, including short copy-number oscillations. Cluster 4 is characterised on average by the longest chains of oscillations and the shortest non-diploid segments, but breakpoint density is moderate. Finally, cluster 5 contains chromosomes with medium/short non-diploid segments, medium/high breakpoint density, and medium/short chains of oscillations.

The distribution of MCF10A chromosomes among clusters is not uniform, with clusters 1 and 2 enriched in MCF10A chromosomes, whereas clusters 3, 4, and 5 are depleted (permutation test, FDR < 0.05). Given that clusters 1 and 2 are characterised by relatively large size copy-number alterations (CNAs) and few breakpoints, we infer that these are likely to represent early stages of chromosome evolution, acquired in one or few cell cycles. Overall, the four clusters that included at least one MCF10A chromosome contained 91% of all analysed PCAWG chromosomes, with the two clusters that are enriched in MCF10A chromosomes containing 43% of all analysed PCAWG chromosomes. The remaining clusters, by comparison, are likely to represent later stages of somatic karyotype evolution in cancer or, in alternative, they could be generated by other processes. In this regard, we note that cluster 4 is characterised by CNAs of less than 200 kb on average – too small to be reliably detectable using Strand-seq[1]. Clusters 3 and 5, by comparison, are characterised by a high number of CN states, indicative of extensive chromosome remodelling across a larger number of cell divisions in combination with Darwinian selection[7] (**Fig. S14D**). In summary, these findings suggest that the CA patterns generated by spontaneous micronucleation broadly reflect those observed in pan-cancer studies, with particular similarity to those chromosome structures that are likely to represent early stages of somatic chromosome evolution in cancer.

### Quantification of aneuploidies in the PCAWG resource

*Quantification of whole chromosome aneuploidies.* To quantify whole-chromosome gains and losses in primary cancer genomes, we leveraged the PCAWG copy number calls[7] (available at https://dcc.icgc.org/api/v1/download?fn=/PCAWG/consensus_cnv). We limited our analysis to chromosomes where the only event was a whole-chromosome gain or loss, and no further copy number alterations (CNAs) were present. To this end, we searched for chromosomes that: (1) had only one copy number segment; and (2) had a copy number value above or below the ploidy of the aliquot[7] (whole-chromosome gain or loss, respectively), regardless of the magnitude of the difference. We defined the ploidy of each aliquot as the most frequent copy number state across all genomic bases that had copy number information. We called all chromosomes that passed this first set of filters our "unfiltered" dataset. Notably, in our unfiltered dataset, 3582/5139 (69.7%) events represent whole-chromosome losses (**Fig. S14A**).

In some aliquots, ploidy could not be unambiguously assigned to an integer number. To exclude these aliquots from the analysis, we filtered out those in which the number of genomic bases affected by the most frequent copy number state was less than three times larger than the number of genomic bases affected by the second most frequent copy number state. In this dataset, 1787/2215 (80.7%) whole-chromosome events are losses (**Fig. S14A**). As an alternative approach, we excluded all PCAWG aliquots that exhibited a whole-genome duplication (WGD) event, as inferred previously[33] for the PCAWG resource. Notably, also in this non-WGD dataset, the vast majority of *de novo* whole chromosome alteration events (1645/2017; 81.5%) represent chromosome losses, with chromosomal gains being comparably rare.

*Analysis of 7q losses in breast cancer*. To assess the frequency of chromosome 7q losses in breast cancer, we reanalyzed copy number data of PCAWG representative aliquots classified as "Breast-AdenoCA" (n = 195), "Breast-LobularCA" (n = 13), or "Breast-DCIS" (n = 3). We considered chromosome 7 to have a q arm loss if >80% of the q arm had a CN value below the inferred ploidy of the aliquot. This restricted the analysis to 78 breast cancer aliquots that had non-ambiguous ploidy, out of which 4 (5.1%) had 7q losses (**Fig. S14F**). Relaxing the filter of % of base positions with lower CN to a more permissive 50% led to only two additional hits (6/78, 7.7%), and they were compatible with terminal losses as well as with extensive chromothripsis.

### Inference of isochromosomes using bulk WGS data

*Methodological approach*. To infer a highly curated set of likely isochromosomes in human cancer cohorts, we designed a restrictive set of criteria that prioritised specificity over sensitivity of detection. We performed the search in both the PCAWG and TCGA datasets, omitting 698 PCAWG donors that were redundant with TCGA (N = 1985 for PCAWG; N = 4957 for TCGA, combined N = 6942) (**Supplementary Table 5**). We excluded sex and acrocentric chromosomes from this analysis. First, we smoothened the CN profiles by removing segments smaller than 10 kb and merging consecutive segments with the same CN state. We excluded chromosomes with more than 150 CAs to filter out highly rearranged chromosomes. Next, we applied the following isochromosome search criteria in forward and reverse directions on each chromosome to find putative isochromosome-like structures in both the p and q-arms. For the detection of isochromosomes of the q-arm, the search was conducted from the beginning of the chromosome to the centromere, while for the p-arm, the search was conducted from the end of the chromosome towards the centromere. First, we searched each chromosome arm for CN changepoints where the minor CN transitions from 0 to a non-zero CN state, only considering changepoints that mapped closer to the centromere than to the telomere. Next, for each CN changepoint, we estimated the modal CN state upstream and downstream of the changepoint for both the minor and the total CN. Finally, we applied the following criteria to identify isochromosome-like structures:

a. The modal CN values, both upstream and downstream of the changepoint, correspond to more than 90% of the sequence.
b. The modal minor CN value is 0 upstream of the changepoint and different from 0 downstream.
c. Downstream of the changepoint, the modal total CN is at least 2 units greater than the modal minor CN.
d. The modal total CN downstream the changepoint is at least 2 units greater than the upstream total CN, and the difference is a multiple of 2.
e. Either (i) the modal total CN upstream of the changepoint is equal to the modal minor CN downstream, or (ii) the sum of the modal total CN upstream of the changepoint and the modal minor CN downstream equals the modal total CN downstream.
f. No more than 2 CAs occur between the changepoint and the centromere, and fewer than 30 CAs occur downstream of the changepoint.

If more than one changepoint satisfied the criteria above for a given chromosome, we selected the changepoint that resulted in the modal CN values with the highest frequencies both upstream and downstream of the changepoint.

*Quantification of centromere-breakpoint distances.* We inferred 1,060 isochromosomes (344 in PCAWG and 716 in TCGA; **Supplementary Table 6**) using this approach. Out of the cohorts that have more than 25 donors, the most frequent isochromosome after normalising by cohort size is i(17q) in PBCA-DE (48/230, 20.9%) (**Fig. S14E**). The PBCA-DE cohort includes paediatric brain cancers such as medulloblastoma, in which highly recurrent i(17q) events have been previously described as highly recurrent[34]. The second most recurrent event is i(8q) in hepatocellular carcinoma (LIRI-JP: 32/250, 12.8%), followed by i(17q) in colon adenocarcinoma (TCGA-COAD: 16/127, 12.6%) and i(8q) in head and neck squamous cancer (TCGA-HNSC: 51/432, 11.8%). Overall, the most frequent isochromosomes across cohorts are i(8q) (n = 238), i(17q) (n = 210), i(3q) (n = 108), and i(5p) (n = 96), all of which have been previously described as highly recurrent[34]. In conclusion, our method, reassuringly, retrieves previously described recurrent isochromosomes.

Encouraged by these observations, we assessed the distances between the inferred isochromosome internal breakpoint and the centromere. The proportion of inferred isochromosomes that have the breakpoint outside the centromere, and thus are likely to be dicentric, is 55.2% in PCAWG and 30.7% in TCGA (**Fig. S14B**). The difference in proportions could be due to differences in assembly (hg19 in PCAWG, hg38 in TCGA), cancer type compositions of each cohort, and data quality, thus making it difficult to compare results between cohorts. It is reassuring to note that for the inferred dicentric isochromosomes, the median distance between the centromere and the internal breakpoint is 3.2 Mb in PCAWG (IQR = 1.9-6.6 Mb) and 2.9 Mb in TCGA (IQR = 1.1-6.1 Mb) (**Fig. S14C**), showing good agreement across the two cohorts.In addition, the distribution of distances extends up to 37 Mb in PCAWG and 42 Mb in TCGA, with inter-centromeric distances of approximately 20 Mb observed commonly. In summary, our results suggest a heterogeneous distribution of isochromosome changepoints, with a substantial proportion of them being megabases away from the centromere.
